# Deficiencies in the Fanconi Anemia or the Homologous Recombination pathway enhance the antitumor effects of the novel hypoxia-activated prodrug CP-506

**DOI:** 10.1101/2025.05.21.655302

**Authors:** Lesley Schuitmaker, Alexander M.A. van der Wiel, Natasja G. Lieuwes, Rianne Biemans, Nikki A.M. Mutsters, Jennifer Jung, Victoria Claudino Bastos, Èlia Prades Sagarra, Sheng Kuang, Sabine A.S. Langie, Jeremy Setton, Jan Theys, Ala Yaromina, Ludwig J. Dubois, Philippe Lambin

**Author notes:** contributed equally to this paper (Lesley Schuitmaker, Alexander M.A. van der Wiel). contributed equally to this paper (Ludwig J. Dubois, Philippe Lambin).

## Abstract

The novel hypoxia-activated prodrug CP-506 has been shown to selectively target hypoxic tumor cells, which are associated with disease progression and resistance to conventional anti-cancer therapies. Given the alkylating effector metabolites, we hypothesize that defects in interstrand crosslink (ICL) and double strand break (DSB) DNA repair may serve as a predictive biomarker of sensitivity to CP-506. Here, we evaluated the role of DNA damage repair pathways in the antitumor response to CP-506.

Isogenic cancer cell lines proficient or deficient in the Fanconi Anemia (FA), homologous recombination (HR), or non-homologous end joining (NHEJ) pathway were cultured as 2D monolayers and 3D spheroids. Cell viability, clonogenic cell survival, and spheroid growth inhibition were assessed following CP-506 exposure. Mice bearing subcutaneous isogenic xenografts received CP-506 (600 mg/kg; QD5) or vehicle treatment upon reaching a tumor starting volume (SV) of 244.9 ± 72.0 mm^3^. Treatment response was quantified as time to reach 4xSV (T4xSV) and respective enhancement ratios (ER), defined as T4xSV_CP-506_/T4xSV_vehicle_. DNA damage and repair capacity were evaluated by γH2AX and alkaline comet assays.

*In vitro*, cell lines deficient in FA or HR, but not NHEJ, showed enhanced sensitivity to CP-506 compared to parental cells in viability and clonogenic assays. This was confirmed in spheroid growth inhibition studies. *In vivo*, the antitumor response to CP-506 was more pronounced (P<0.0001) in LNCaP AR *FANCA^-/-^* (ER 4.0±1.1) and LNCaP AR *FANCD2^-/-^* (ER 3.4±0.8) xenografts compared to parental LNCaP AR (ER 1.5±0.5) xenografts. CP-506 treatment of DLD-1 *BRCA2^-/-^* (2.9±0.7; P<0.0001) xenografts resulted in significantly enhanced ER compared to parental (1.3±0.2) xenografts. Similar results were obtained in HCT116 *BRCA2^-/-^* (ER 4.0±0.6; P<0.0001) versus parental (ER 1.7±0.6) xenografts. In contrast, the ER of HCT116 DNA-PKcs^-/-^ (1.4±0.3; P=0.18) xenografts was not different from HCT116 parental xenografts. Under anoxic conditions, CP-506 caused elevated γH2AX foci counts in FANCA-(2.0-fold increase at 48 hours) and FANCD2-deficient cells (1.4-fold increase at 72 hours) compared to LNCaP AR parental cells (P<0.0001). Similarly, γH2AX expression was increased in DLD-1 *BRCA2^-/-^* cells, but not in HCT116 *BRCA2^-/-^*cells, compared to their respective parental cells. In xenografts, FA- and HR-deficiency caused elevated γH2AX expression compared to respective parental tumors (1.6-9.3-fold increase). HCT116 *DNA-PKcs^-/-^* cells and xenografts displayed reduced γH2AX expression compared to their parental counterpart, with an 0.5-fold reduction. The comet assay confirmed CP-506–induced ICLs and DNA strand breaks but was unable to explain the differential therapeutic responses of CP-506 among isogenic tumor cells.

Deficiencies within FA or HR, but not NHEJ, enhanced the antitumor effects of CP-506 through a mechanism consistent with the concept of synthetic lethality. Therefore, both DNA repair status and the presence of tumor hypoxia represent key biomarkers for patient stratification in clinical trials of CP-506.

## Introduction

Hypoxia-activated prodrugs (HAPs) are a class of cytotoxic agents that selectively target and eliminate hypoxic tumor cells, which are associated with disease progression (1) and resistance to conventional anti-cancer therapies (2). Several HAPs have been evaluated in both preclinical and clinical settings (3, 4). Despite highly encouraging preclinical results, implementation of HAPs in the clinic has not been successful to date, which can be, at least in part, attributed due to a lack of patient stratification in the design of these clinical trials (5). The identification of key factors influencing the tumoral response to HAPs and respective biomarkers of response are therefore essential for successful clinical implementation.

At least three factors are proposed to influence the antitumor effects of a HAP (5): first, the degree and severity of tumor hypoxia; second, the levels and activity of endogenous oxidoreductases effecting an initial activation step to yield an oxygen-sensing intermediate; and third, the intrinsic sensitivity of the tumor cell to the effector molecules of the HAP (6).

Recently, we have validated the hypoxia-selectivity of CP-506, a novel hypoxia-activated DNA crosslinking prodrug (7). CP-506 was selectively activated only under severe hypoxic (< 0.1% O_2_) conditions *in vitro* and demonstrated potent antitumor effects *in vivo* in a broad range of hypoxic tumor xenograft models. Furthermore, a causal relationship between tumor oxygenation and its therapeutic efficacy was established. Moreover, we identified the one-electron reductases cytochrome P450 oxidoreductase (POR), methionine synthase reductase (MTRR), novel diflavin oxidoreductase 1 (NDOR1), and inducible nitric oxide synthase (NOS2A) as likely candidates for the required initial activation step of CP-506 (7).

As bifunctional alkylators, the hydroxylamine and amine effector metabolites of CP-506 induce various forms of DNA damage, including interstrand crosslinks (ICLs) and monoadducts, as evidenced by the hypoxia-selective formation of DNA adducts (7, 8). ICLs are extremely toxic DNA lesions because the covalent linkage between the two DNA strands prevents DNA strand separation and thereby interferes with DNA replication and transcription. Repair of ICLs involves a complex and highly coordinated response of components of the Fanconi anemia (FA) pathway or nucleotide excision repair (NER) (9). ICL repair proceeds via the formation of double-strand DNA break (DSB) intermediates with subsequent error-free repair by homologous recombination (HR) in the S-phase or by error-prone repair by non-homologous end joining (NHEJ) in all phases of the cell cycle (10-12).

The intrinsic sensitivity of tumor cells to the effector molecules of CP-506 is therefore likely determined by the integrity of the DNA damage response (DDR), able to recognize and repair the induced DNA damage (13). Supportive of this, we recently discovered that the MDA-MB-468 cell line – most responsive to CP-506 treatment in a panel of 15 different *in vivo* xenograft models (7) – is defective in the FA pathway, harboring a truncating mutation (Q869*) in the FA complementation group A (*FANCA*) gene. In addition, several studies have demonstrated that cells and tumors deficient in HR are more sensitive to DSB- and crosslink-inducing chemotherapies (14-16), poly (ADP-ribose) polymerase inhibitors (PARPi) (17, 18), and PR-104 and evofosfamide (TH-302) (19-21), HAPs with a similar mechanism of action as CP-506.

The present study aimed to evaluate the role of different DNA repair pathways in determining the antitumor efficacy of CP-506. We hypothesized that cancer cells and tumors deficient in DNA repair pathways involved in repair of ICLs and DSBs may exhibit increased sensitivity to CP-506, consistent with the concept of synthetic lethality. *In vitro*, we first determined the sensitivity of isogenic cancer cell lines, proficient or deficient in the FA, HR, or NHEJ pathway, to CP-506 using cell viability assays and clonogenic cell survival assays, after which we validated our findings in 3D spheroid models. *In vivo*, we further characterized the role of these DNA repair pathways in the antitumor effects of CP-506 in isogenic xenograft models. Lastly, DNA damage and repair were assessed by γH2AX – a marker of DSB – expression (22) and by alkaline comet assays (23, 24).

## Materials and methods

### Compounds

CP-506 (2-[(2-bromoethyl)-5-[(4-ethyl-1-piperazinyl)carbonyl]-2-(methylsulfonyl)-4-nitroanilino]ethyl methanesulfonate) was manufactured by Mercachem employing synthetic routes developed at the University of Auckland (25). Chlorambucil (4-[bis(2-chloroethyl)amino]benzenebutyric acid, 4-(4-[bis(2-chloroethyl)amino]phenyl)butyric acid) was purchased from Sigma Aldrich. For *in vitro* experiments, compounds were prepared in dimethyl sulfoxide (DMSO) and stored at -20 °C. For *in vivo* experiments, CP-506 was dissolved in water for injection (WFI) and chlorambucil in 48% PEG-400 in WFI (v/v).

### Cell culture

Cells were cultured at 37°C in a humidified 5% CO_2_ air atmosphere and were short tandem repeat (STR)-authenticated and confirmed to be mycoplasma-free by using the MycoAlert™ Mycoplasma Detection Kit (Lonza). Tissue of origin, genetic mutation and corresponding DNA repair pathway affected, provider, and culture medium are reported in Supplementary Table S1. Culture medium for all 2D assays was preincubated 24 hours before use in normoxic or anoxic conditions in a cell culture incubator (HERAcell® 150 CO_2_ Incubator; 21% O_2_, 5% CO_2_) or an anoxic workstation (A35 Don Whitley, Don Whitley Scientific; < 1 ppm O_2,_ 10% H_2_, 5% CO_2_, residual N_2_). Upon overnight attachment, cells were transferred to normoxic or anoxic conditions and received preincubated culture medium. After 24 hours, cells were exposed to CP-506-containing preincubated medium for 4 hours.

### Cell viability assays

Cells were seeded in 96-well plates in optimized cell densities. After CP-506 treatment, plates were transferred to normoxic conditions, washed with phosphate-buffered saline (PBS), and received fresh culture medium. Cell viability was assessed 72 hours after the start of treatment using the alamarBlue™ reagent according to the manufacturer’s instructions. Treatment response was quantified as IC_50_, i.e. the concentration of CP-506 that resulted in a 50% reduction in cell viability.

### Clonogenic cell survival assays

Cells were seeded in 60-mm glass dishes in optimized cell densities. After CP-506 incubation, cells were transferred to normoxic conditions, washed, harvested, and seeded as single cell suspensions to assess clonogenic cell survival after ∼12 days. Colonies (> 50 cells) were manually counted to determine plating efficiency, after which survival fractions were calculated.

### Spheroid culture

Spheroids were grown as described previously (26). Spheroid growth was monitored using an IX81 inverted microscope (Olympus), equipped with an ORCA-Fusion C14440 20-UP camera (Hamamatsu), using the µManager open-source microscopy software (27). Spheroid volume was determined using the MATLAB-based open-source SpheroidSizer software (28). Once spheroids were hypoxic, i.e. reaching a spheroid volume of ca. 0.20 mm^3^ as determined by pimonidazole positivity previously (7), spheroids were treated with CP-506 for 24 hours. Afterwards, spheroids were washed and received fresh culture medium. Treatment response was quantified as spheroid growth inhibition (SGI) 7 days post start of treatment, calculated as 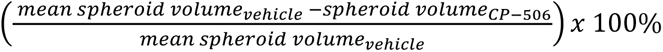.

### Animals

All animal experiments were performed with appropriate ethical approval by the Centrale Commissie Dierproeven (AVD1070020198905), according to institutional guidelines of Maastricht University. Mouse strains are specified in Supplementary Table S2.

#### Therapeutic response study

Cells (Supplementary Table S1) were resuspended in Matrigel™ (BD Biosciences) and injected subcutaneously into the flank of the animal. Upon reaching a tumor volume of 244.9 ± 72.0 mm^3^ (treatment starting volume, SV), mice were randomly assigned to following treatment groups: vehicle (WFI), chlorambucil (3 mg/kg), or CP-506 (600 mg/kg) for five consecutive days (QD5, IP). Tumor growth was assessed by measuring the tumor in three dimensions using a Vernier caliper and using the formula 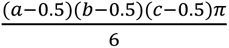 where a, b, and c are orthogonal diameters of the tumor and 0.5 mm is a correction for the thickness of the skin. Tumor response was quantified as (i) tumor growth inhibition (TGI), defined as 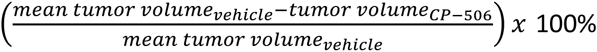, the day respective control animals reached four times SV (4xSV) and (ii) as the enhancement ratio (ER) defined as the ratio of time to reach 4xSV (T4xSV) of CP-506 treated animals to T4xSV of vehicle-treated animals.

#### Histology study

Isogenic xenograft-bearing mice were randomized to assess DNA damage at 48 hours post start of treatment. Upon reaching a tumor volume of 223.9 ± 30.1 mm^3^, mice were treated (QD1, IP) with vehicle (WFI) or CP-506 (600 mg/kg). Pimonidazole (60 mg/kg dissolved in 0.9% saline, IP) was injected one hour before tumor excision. Half of a tumor was fixed in 4% (v/v) formalin and embedded in paraffin for the detection of γH2AX, the other half was snap-frozen in liquid nitrogen and stored at - 80°C for the detection of pimonidazole to assess the hypoxic fraction (HF) as described previously (29) (Supplementary Methods).

### Assessment of DNA damage

#### In vitro γH2AX immunofluorescence

Cells were seeded in 35-mm glass dishes in optimized cell densities. Isogenic cancer cell lines were treated with anoxic IC_50_ values of respective parental cell lines (LNCaP AR: 73.2 µM, DLD-1: 158.6 µM, HCT116: 65.2 µM). After CP-506 treatment under anoxic conditions, cells were transferred to normoxic conditions, washed, and received fresh culture medium. For immunofluorescence detection of γH2AX, cells were fixed 48 and 72 hours post start of treatment using methanol (-20 °C) for 15 min. Thereafter, cells were washed and permeabilized in 0.2% (v/v) Triton X-100 (Thermo Fisher Scientific) in PBS for 10 min at room temperature (RT). Next, non-specific binding was blocked using 5% (v/v) normal goat serum (Thermo Fisher Scientific) in 0.02% (v/v) Triton X-100 in PBS for 20 min at RT. Cells were incubated with primary anti-phospho-H2A.X (Ser139) antibody (clone JBW301, Merck, 1:500) for 2 hours, after which cells were washed and incubated with Alexa Fluor® 488-conjugated goat anti-rabbit IgG antibody (1:500; Invitrogen) for 1 hour all in a humidified box at 37°C. Cells were washed and nuclei were stained using Hoechst (1:5000; Thermo Fisher Scientific) for 10 min at RT. Slides were imaged using a Leica DMI 4000 confocal microscope (Leica) using a 60x oil immersion objective.

Image fluorescence was quantified using ImageJ software version 1.54f (National Institutes of Health) (30) in a semi-automated manner. First, maximum intensity projections of Z-stacks were made to construct 2D images. For each constructed image, single channel Hoechst images were used to determine nuclei as regions of interest. Next, to differentiate between the foci and background, manual thresholds were set by two independent researchers (N.A.M.M. and L.S.). Finally, γH2AX foci per nucleus were counted for LNCaP AR images and γH2AX fluorescence intensity per nucleus was quantified for HCT116 and DLD-1 images.

#### Ex vivo γH2AX immunohistochemistry

Paraffin-embedded tumor material from isogenic xenografts was sectioned to assess γH2AX expression 48 hours post start of treatment, as previously described (29). Tumor sections (7 μm) were deparaffinized (xylene, 20 min) and rehydrated using a graded ethanol series. For antigen retrieval, slides were microwaved in sodium citrate buffer solution for 20 min and cooled on ice. Endogenous peroxidases blocking was performed using 3% peroxidase solution (5 min), after which slides were washed in tris-buffered saline (TBS) with 0.2% (v/v) Tween (Merck, TBS-Tw). To prevent unspecific antibody binding, slides were incubated for 30 min with 3% (w/v) bovine serum albumin (Carl Roth) in TBS-Tw. Slides were incubated with a mouse monoclonal anti-phospho-Histone H2A.X (Ser139) antibody (clone JBW301, biotin conjugate, Merck, 1:250) overnight at 4°C, followed by incubation with the VECTASTAIN Elite ABC Kit reagents (Vector Laboratories) according to the manufacturer’s instructions. As chromogen, 3’3’-Diaminobenzidin (Sigma-Aldrich) was used and counterstaining was performed using hematoxylin (Klinipath). Finally, the slides were mounted with coverslips using DPX mounting medium (Brunschwig Chemie).

Tumor sections were imaged using a Precipoint M8 microscope and scanner equipped with a 20x objective. For the quantification of γH2AX staining, ImageJ version 1.54f and QuPath version 0.4.3 (31) were used. First, vital tumor regions, excluding necrotic areas, connective tissue, and processing or staining artefacts, were segmented in an automated manner using a deep-learning DynUNet model (Supplementary Methods). If necessary, manual corrections of the automated vital masks were performed in ImageJ, after which the vital masks were overlaid with the original image. Overlaid images, solely containing the vital tumor regions, and an original image per isogenic tumor model were imported in QuPath (Supplementary Figure S1). The staining vectors were set on the original image, after which the tissue boundaries of the overlaid images were detected by means of the simple tissue detection function, followed by the detection of individual cells and nuclei using the positive cell detection function with optimized parameters per isogenic tumor model (Supplementary Table S3). To define γH2AX positive nuclei, the thresholds were set manually per isogenic tumor model according to signal intensity and background staining by one investigator (L.S.) blinded to subject coding. The percentage of γH2AX positive nuclei was calculated as the number of nuclei with γH2AX intensity above the set threshold divided by the total amount of nuclei detected in the vital region of the tumor section. Additionally, an object classifier was trained on representative images of the DLD-1 isogenic model with dedicated annotations to differentiate between tumor regions, necrotic regions, and connective tissue that were not detected by the machine-learning model and too demanding to exclude manually.

### Alkaline comet assay for detection of interstrand crosslinks and DNA strand breaks

To assess single-strand break (SSB) and DSB DNA damage, the standard alkaline comet assay was employed (23). In parallel, detection of ICL was performed using a modified alkaline comet assay, which can detect ICLs by challenging the cells in the gels with H_2_O_2_ as described previously (24).

Isogenic cancer cells were treated under anoxic conditions with CP-506 (50 μM for HCT116; 100 μM for DLD-1 and LNCaP AR). Concentrations were selected based on preliminary dose-response experiments, corresponding to those at which ICLs and DNA damage were detectable using the (modified) alkaline comet assay. CP-506 and vehicle-treated cells were harvested 48 and 72 hours post start of treatment and slowly frozen to -80°C in freezing medium (50% FBS, 45% culture medium, 5% DMSO) until the (modified) alkaline comet assay was performed.

Single-cell suspensions (in cold PBS) were mixed with low melting point agarose at 37°C to a final concentration of 0.7% with 5×10^4^ cells/mL. Then, droplets (7 μL) of the cell-suspension-agarose mixtures were carefully placed on normal melting point agarose pre-coated microscope slides in duplicate, with a total of 12 mini-gels per slide. Once the mini-gels were set (2 min on cold plate), half of the slides were exposed to 100 mM hydrogen peroxide in ice cold PBS for 5 min for the modified alkaline comet assay. After a cold PBS wash, all slides were exposed to lysis solution (2.5 M NaCl, 0.1 M Na_2_EDTA, 10 mM Trizma® base, pH 10, and 1% (v/v) Triton® X-100) for 1 hour at 4°C. Slides were transferred to the electrophoresis tank and immersed in electrophoresis solution (0.3 M NaOH, 1 mM EDTA-Na_2_) for 40 min at 4°C for DNA unwinding. Electrophoresis was performed at 0.94 V/cm for 20 min at 4°C. Afterwards, slides were washed in PBS and MilliQ for neutralization. The mini-gels were then dehydrated with 70% and 100% ethanol for 5 min each. After air-drying, the mini-gels were stained with 100 µl of 3x GelRed stain (Millipore) and covered using a coverslip for visualization. The fluorescence microscope Cytation III (BioTek, Agilent) equipped with a 10x objective was used to acquire images. The comets were analyzed using the semi-automated image analysis software Comet Assay IV (Instem Perceptive Instruments). All analyses were performed by one investigator (R.B.) blinded to experimental labeling. Tail intensity was quantified as a percentage of DNA in tail (%DNA in tail), i.e. the pixel intensity of the tail respective to the total DNA content. 50 random nuclei per mini-gel, 100 per experimental condition, were scored.

### Statistics

Statistical analyses were performed using GraphPad Prism 10.1.2 software (GraphPad Software, Inc.). Cell viability and clonogenic cell survival curves were fitted to an inhibitory dose-response curve and linear quadratic model as a function of CP-506 concentration, respectively, after which the parameters of the curves were compared between the isogenic cancer cell lines with their respective parental control. Two-sided t-test or one-way ANOVA with Dunnett’s multiple comparison test were performed to assess statistical significance between curve fit parameters. A two-way ANOVA was used to evaluate statistical significance in SGI, TGI, ER, T4xSV, percentage of γH2AX positive cells, and %DNA in tail, the means of the parental and respective isogenic xenografts or cells for the different treatment arms followed by the Dunnet’s or Tukey’s multiple comparisons test. A non-parametric Kruskal Wallis test with Dunn’s multiple comparison was performed to test differences in γH2AX immunofluorescence staining. Data are reported as mean ± standard deviation (SD) or median (IQR). Results were considered significant if the p-value was < 0.05 (*), < 0.01 (**), < 0.001 (***), or < 0.0001 (****).

## Results

Cells deficient in the Fanconi anemia or homologous recombination pathway are more sensitive to CP-506 To investigate the role of DNA repair pathways on the efficacy of CP-506, we first assessed cell viability of the different isogenic cancer cell lines under normoxic and anoxic conditions upon increasing concentrations of CP-506. For all isogenic cell lines tested, normoxic IC_50_ values were consistently higher than anoxic IC_50_ values resulting in hypoxia-cytotoxicity ratios (HCRs) ranging from 3.2 to 20.3, supportive of the hypoxia-dependent metabolism and cytotoxicity of CP-506. Deficiencies in FA resulted in higher sensitivity to CP-506 compared to parental cells (Table 1). Similarly, isogenic cancer cells deficient in HR were more sensitive to CP-506 under anoxic conditions compared to their respective parental controls. The isogenic cancer cells deficient in NHEJ (*DNA-PKcs*^-/-^) were less sensitive to CP-506 under anoxic conditions, as indicated by higher anoxic IC_50_ values compared to their respective parental control (Table 1). Complete dose-response curves under normoxic and anoxic conditions are shown in Supplementary figure S2.

**Table 1.** IC50 values of *in vitro* monolayer cultures of isogenic cancer cell lines proficient or deficient in DNA repair pathways after exposure to increasing concentrations of CP-506 under normoxic (21% O2) or anoxic (≤ 0.02% O2) conditions.

| Cell line | Cancer type | DNA repair pathway | NIC<sub>50</sub> | AIC<sub>50</sub> | HCR | P-value |
| --- | --- | --- | --- | --- | --- | --- |
| LNCaP AR | Prostate | Parental | 471.5 | 73.2 | 6.4 | - |
| LNCaP AR *FANCA*<sup>-/-</sup> | Prostate | FA | >500.0 | 36.4 | >13.7 | ns |
| LNCaP AR *FANCD2*<sup>-/-</sup> | Prostate | FA | 302.0 | 25.7 | 11.8 | ns |
| DLD-1 | Colorectal | Parental | >500.0 | 158.6 | >3.2 | - |
| DLD-1 *BRCA2*<sup>-/-</sup> | Colorectal | HR | >500.0 | 72.5 | >6.9 | <0.0001 |
| HCT116 | Colorectal | Parental | >500.0 | 65.2 | >7.7 | - |
| HCT116 *BRCA2*<sup>-/-</sup> | Colorectal | HR | 387.8 | 19.1 | 20.3 | <0.001 |
| HCT116 *DNA-PKcs*<sup>-/-</sup> | Colorectal | NHEJ | >500.0 | 103.9 | >4.8 | ns |

NIC<sub>50</sub>: normoxic IC<sub>50</sub> value; AIC<sub>50</sub>: anoxic IC<sub>50</sub> value; HCR: hypoxia-cytotoxicity ratio. P-values are determined by comparison of curve fit parameters between the isogenic cancer cell lines and their respective parental cell lines.

These findings were validated using clonogenic cell survival assays. Under normoxic conditions, CP-506 exposure only marginally affected clonogenic cell survival in any of the isogenic cancer cell lines tested (Supplementary Figure S3). Under anoxic conditions, clonogenic survival decreased with increasing CP-506 concentrations (Figure 1A-C). Compared to their respective parental controls, clonogenic cell survival was significantly decreased in LNCaP AR cancer cells deficient in FANCA (P < 0.05) and FANCD2 (P < 0.001; Figure 1A). Similarly, deficiencies in BRCA2, but not in DNA-PKcs, sensitized HCT116 (P < 0.01) and DLD-1 cancer cells (P < 0.0001) to CP-506 treatment (Figure 1B and 1C).

**Figure 1.**
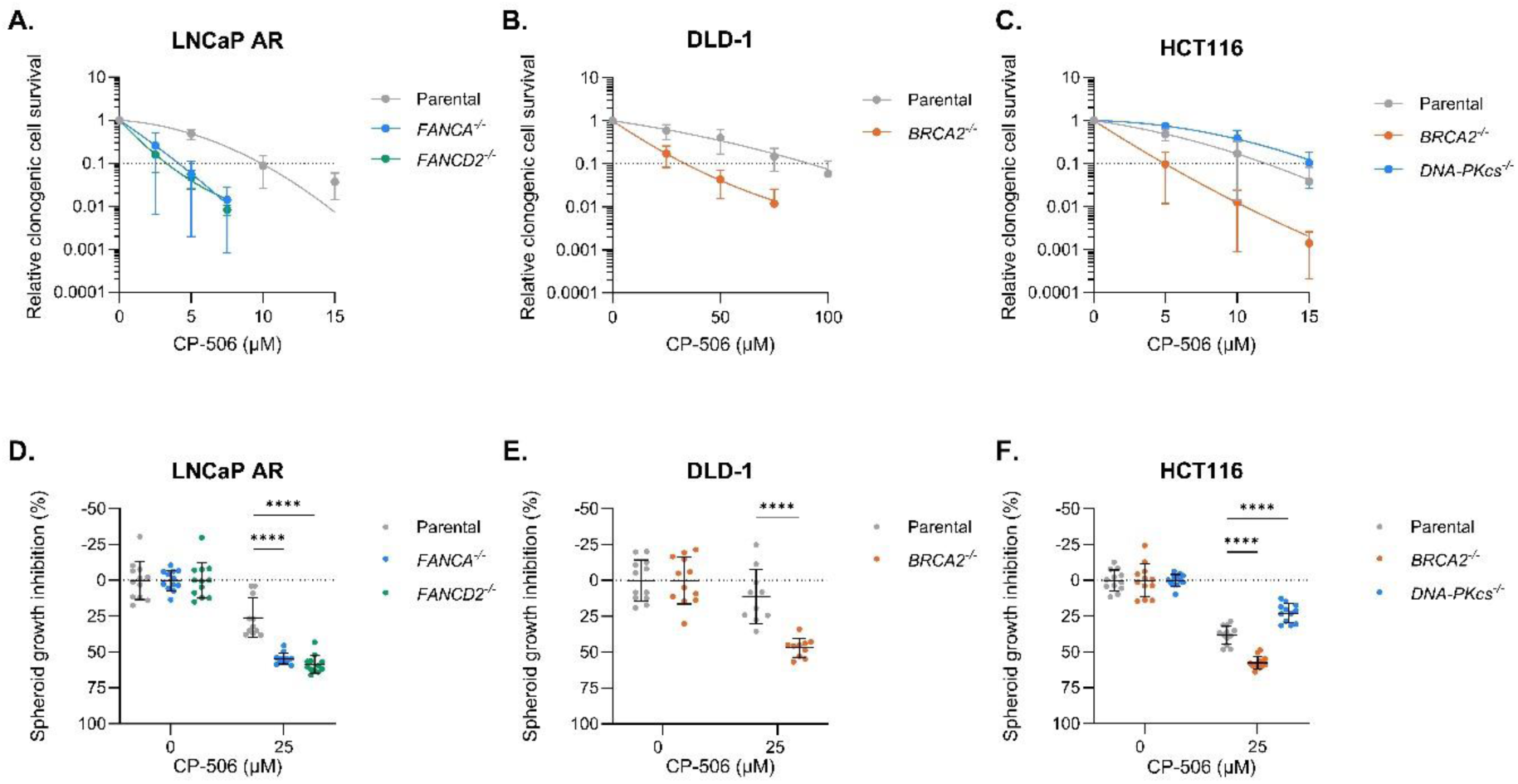
Deficiencies in the Fanconi anemia or the homologous recombination pathway sensitize cancer cells and spheroids to CP-506. Clonogenic cell survival after 4 hours of anoxic exposure to CP-506 of LNCaP AR (A), DLD-1 (B), and HCT116 (C) isogenic cell lines proficient or deficient in FA, HR, or NHEJ. Spheroids were treated with CP-506 for 24 hours and spheroid growth inhibition was assessed 7 days post start of treatment of LNCaP AR (D), DLD-1 (E), and HCT116 (F) isogenic spheroids proficient or deficient in FA, HR, or NHEJ. Per condition, 10-12 spheroids were used. Data are presented as mean ± SD of ≥ 3 independent experiments. ****: P < 0.0001.

Hypoxic 3D spheroid cultures were used to further confirm the role of DNA repair pathways in the cytotoxicity of CP-506. In all spheroid cultures, CP-506 induced an SGI at 7 days post start of treatment. FA-deficient LNCaP AR spheroids were significantly more sensitive to CP-506 as indicated by a more pronounced SGI in FANCA-(54.5 ± 3.8%; P < 0.0001) and in FANCD2-deficient spheroids (58.4 ± 6.2%; P < 0.0001) when compared to parental LNCaP AR spheroids (26.0 ± 13.7%) (Figure 1D). In DLD-1 spheroids, CP-506 exposure induced a stronger SGI in spheroids deficient in BRCA2 (46.9 ± 6.6%) compared to parental spheroids (11.2 ± 18.8%; P < 0.0001) (Figure 1E). In HCT116 spheroids, BRCA2-deficiency also resulted in a stronger SGI (57.4 ± 4.3%) compared to parental spheroids (38.1 ± 6.4%; P < 0.0001). In line with the 2D *in vitro* results, spheroids deficient in DNA-PKcs were significantly less sensitive (SGI: 22.8 ± 6.5%; P < 0.0001) to CP-506 treatment as compared to parental HCT116 spheroids (Figure 1F). These data further confirm that cells and spheroids deficient in FA and HR, but not in NHEJ, exhibit increased sensitivity to CP-506.

The antitumor effects of CP-506 are enhanced in xenografts deficient in the Fanconi anemia or the homologous recombination pathway To further evaluate the role of DNA repair pathways in the single-agent antitumor activity of CP-506 *in vivo*, mice bearing subcutaneous isogenic xenografts were treated with CP-506. As a functional validation of DNA repair deficiency, mice were treated with the non-hypoxia-activated alkylating agent chlorambucil. In all models tested, CP-506 and chlorambucil were well-tolerated with only transient body weight loss during the treatment period (Supplementary Figure S4). Tumor hypoxia – an essential factor for CP-506 activation – was confirmed in all models tested as determined by pimonidazole positivity (Supplementary Figure S5).

Treatment with CP-506 resulted in a TGI for LNCaP AR parental (52.3 ± 56.4%; P < 0.05), *FANCA^-/-^* (98.6 ± 1.5%; P < 0.0001), and *FANCD2^-/-^* tumors (97.7 ± 3.2%; P < 0.0001; Figure 2A). CP-506 increased T4xSV compared to vehicle-treated controls in LNCaP AR parental (P = 0.29), LNCaP AR *FANCA^-/-^* (P < 0.0001), and LNCaP AR *FANCD2^-/-^* (P < 0.0001). The resulting ERs were significantly higher for LNCaP AR *FANCA^-/-^* (ER: 4.0 ± 1.1; P < 0.0001) and LNCaP AR *FANCD2^-/-^* (3.4 ± 0.8; P < 0.0001) compared to LNCaP AR parental (ER: 1.5 ± 0.5) xenografts (Figure 2B and Supplementary table S2). FA-deficient LNCaP AR xenografts were also significantly more sensitive to chlorambucil (Supplementary Figure S6).

**Figure 2.**
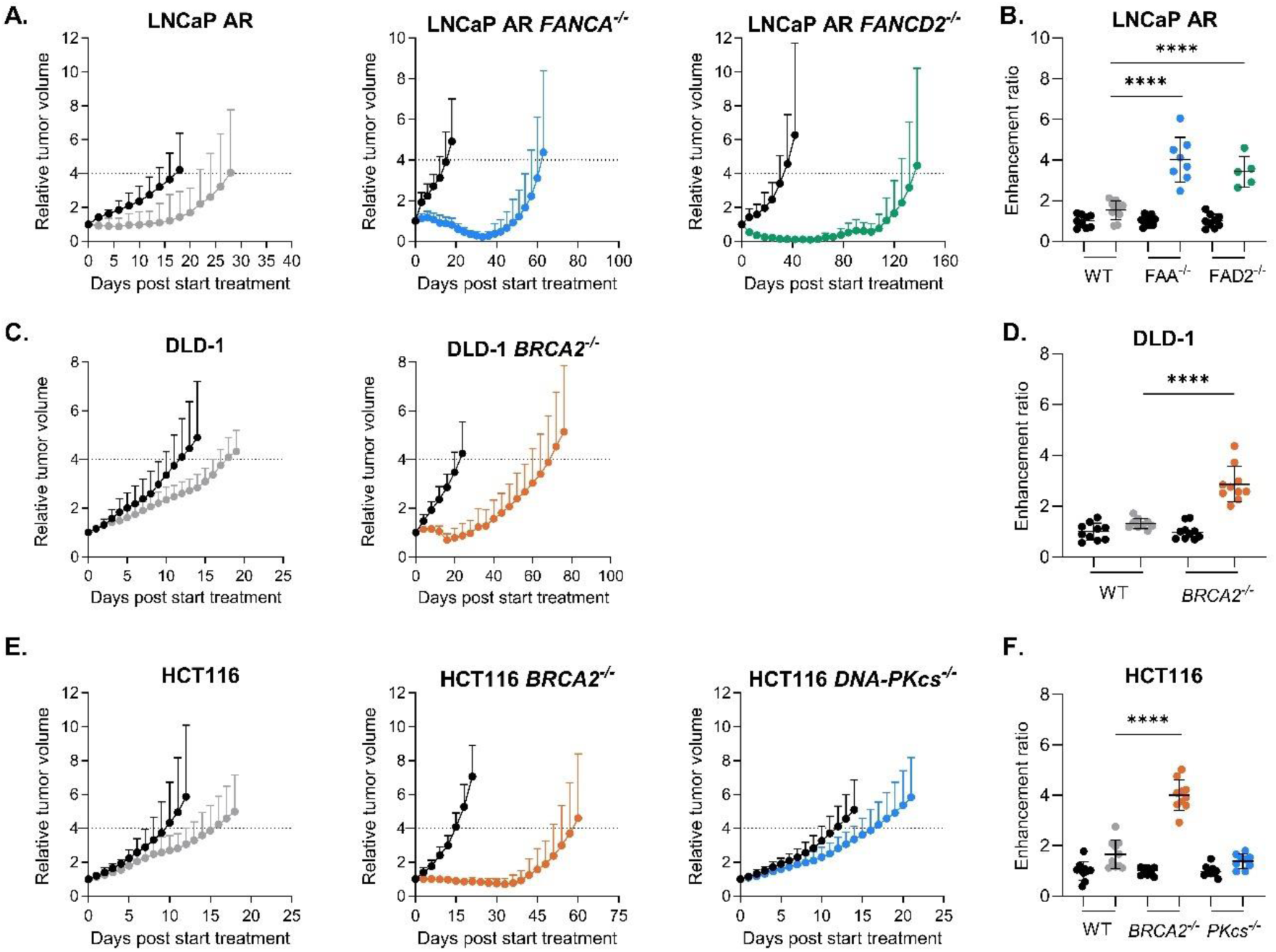
The antitumor effects of CP-506 are enhanced in tumors deficient in the Fanconi anemia or homologous recombination pathway. Mice bearing isogenic tumor xenografts, proficient or deficient in FA (*FANCA*^-/-^ and *FANCD2*^-/-^), HR (*BRCA2*^-/-^), or NHEJ (*DNA-PKcs*^-/-^) were treated with vehicle (black circles) or CP-506 (colored circles) after which tumor growth was monitored (A, C, E) and time to reach four times starting volume (T4xSV) and corresponding enhancement ratio (ER) were determined (B, D, F). Data are presented as mean ± SD (n = 5-10 animals per group). ****: P < 0.0001.

CP-506 effectively inhibited tumor growth when compared to vehicle treatment in DLD-1 parental (37.9 ± 10.3%; P < 0.01) and DLD-1 *BRCA2^-/-^* xenografts (77.8 ± 11.0%; P < 0.0001; Figure 2C). For DLD-1 parental xenografts the T4xSV was not significantly different between vehicle and CP-506-treated animals (P = 0.49), whereas CP-506-treatment significantly prolonged T4xSV in BRCA2-deficient xenografts (P < 0.0001; Supplementary Table S2). Treatment of DLD-1 *BRCA2^-/-^* with CP-506 resulted in a significantly higher ER (2.9 ± 0.7; P < 0.0001) when compared to parental DLD-1 tumors (ER: 1.3 ± 0.2; Figure 2D). Similar responses were observed in HCT116 xenografts. BRCA2-deficient xenografts showed significantly enlarged ERs (4.0 ± 0.6; P < 0.0001) compared to HCT116 parental xenografts (1.7 ± 0.6; Figure 2E and 2F). In HCT116 *DNA-PKcs^-/-^* xenografts, however, CP-506 induced a TGI of 33.2 ± 17.3% (P = 0.15) and significantly increased T4xSV (17.5 ± 3.6) compared to vehicle-treated controls (12.6 ± 2.7; P < 0.05; Figure 2E and Supplementary Table S2). However, the ER of HCT116 *DNA-PKcs^-/-^*xenografts (1.4 ± 0.3; P = 0.18) was not significantly different from HCT116 parental xenografts (Figure 2F and Supplementary table S2). In line with these findings, the antitumor effects of chlorambucil were also more pronounced in HCT116 and DLD-1 xenografts deficient in HR (Supplementary Figure S6). Taken together, these data demonstrate that *in vivo* tumors deficient in FA or HR, but not NHEJ, are more sensitive to CP-506.

### CP-506 induces phosphorylation of histone H2AX

To explore the extent of residual DNA damage after CP-506 exposure in more detail, γH2AX expression was assessed using immunofluorescence in *in vitro* isogenic cancer cells and immunohistochemistry in *ex vivo* isogenic xenografts. Upon anoxic exposure to CP-506, γH2AX expression presented as distinct nuclear foci in isogenic LNCaP AR cells (Figure 3A). 48 hours after CP-506 exposure, all isogenic LNCaP AR cells showed elevated foci counts compared to vehicle treatment (P < 0.0001). FANCA-(55.0 (IQR = 28.0 – 84.0); P < 0.0001) and FANCD2-deficient (37.0 (IQR = 9.5 – 76.8); P = 0.06) LNCaP AR cells expressed higher number of nuclear foci compared to LNCaP AR parental cells (28.0 (IQR = 11.5 – 53.0)). The number of foci per nucleus remained higher in FANCD2-deficient cells 72 hours post treatment (50.0 (IQR = 32.0 – 89.0); P < 0.0001), whereas γH2AX foci in FANCA-deficient cells (34.5 (IQR = 16.0 – 59.8); P = 0.99) reverted to parental level (36.0 (IQR = 10.0 – 52.0); Supplementary Figure S7A).

**Figure 3.**
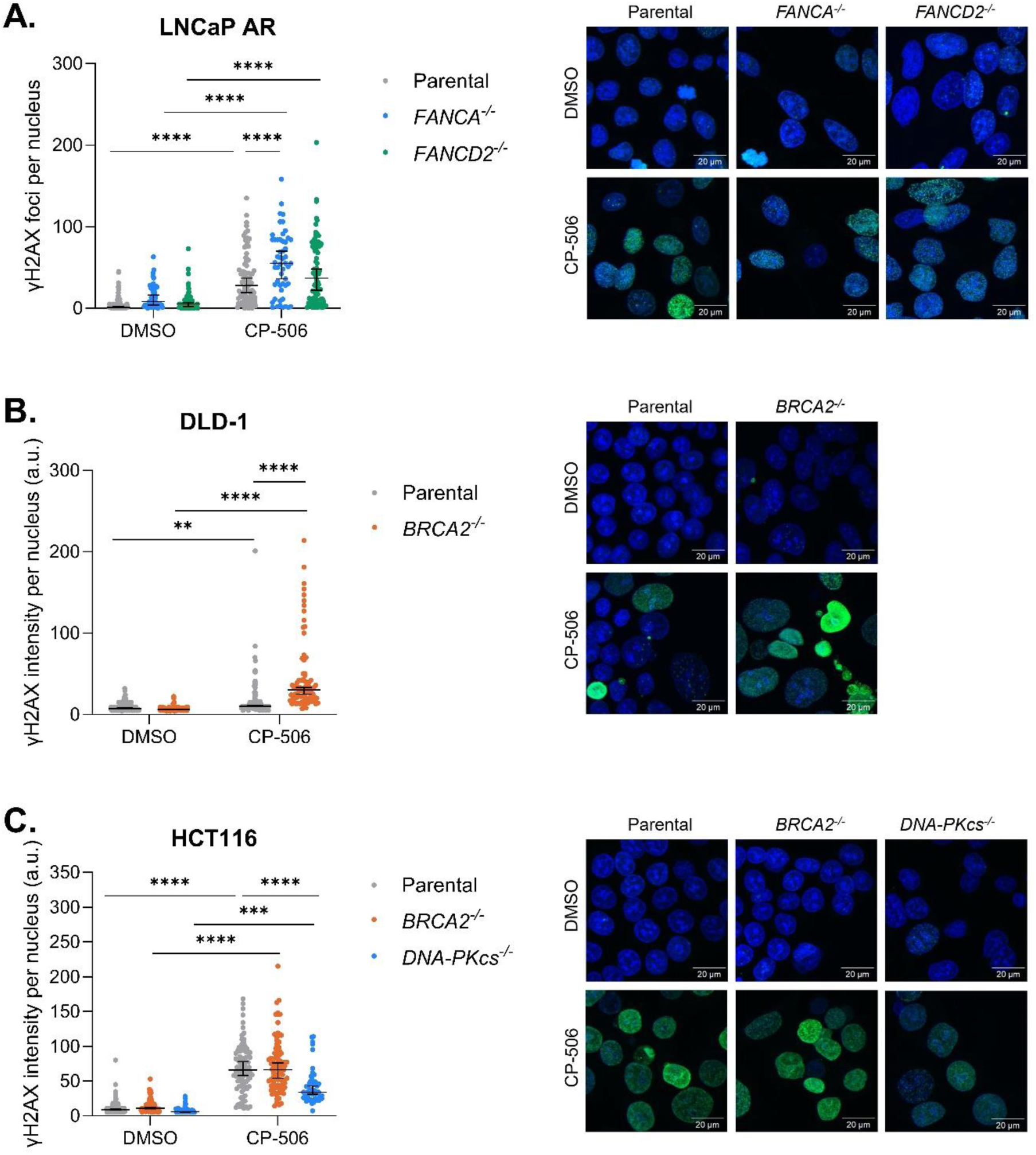
CP-506 induced persistent DNA damage in FA- or HR-deficient isogenic cancer cells 48 hours post start of treatment under anoxic conditions. γH2AX foci count per nucleus for LNCaP AR isogenic cells (A) and quantification of γH2AX immunofluorescence intensity per nucleus for DLD-1 (B) and HCT116 (C) isogenic cancer cells with representative images. Blue: Hoechst; green: γH2AX. n ≥ 42 cells per condition. Data are presented as median (IQR). **: P < 0.01, ***: P < 0.001, ****: P < 0.0001.

Interestingly, in isogenic DLD-1 and HCT116 cells, γH2AX expression presented as a pan-nuclear staining after anoxic exposure to CP-506 (Figure 3B and 3C). Therefore, instead of counting distinct foci, γH2AX fluorescence intensity per nucleus was quantified. Exposure to CP-506 resulted in an elevated (P < 0.0001) expression of γH2AX in DLD-1 BRCA2-deficient cells when compared to DLD-1 parental cells (Figure 3B). The differences in γH2AX expression levels between parental and BRCA2-deficient DLD-1 cells remained at 72 hours after treatment (P < 0.01; Supplementary Figure S7B). In HCT116 cells, BRCA2 deficiency did not result in elevated γH2AX expression levels neither after 48 hours (P = 0.53; Figure 3C), nor after 72 hours post treatment (P = 0.61; Supplementary Figure S7C). HCT116 *DNA-PKcs^-/-^*cells exhibited a significant decrease in H2AX phosphorylation at 48 (P < 0.001; Figure 3C) and 72 (P < 0.0001; Supplementary Figure S7C) hours post treatment compared to their respective parental cancer cells.

Next, γH2AX positivity was assessed in isogenic tumors excised 48 hours post treatment. For LNCaP AR parental and FANCD2-deficient tumors (Figure 4A), the percentage of γH2AX positive cells after CP-506 treatment (3.8 ± 1.9% and 19.3 ± 11.7%, respectively) was similar compared to vehicle-treated tumors (6.1 ± 4.8%; P = 0.78 and 10.4 ± 5.2%; P = 0.24). In contrast, in FANCA-deficient tumors, the percentage of γH2AX positive cells was increased after exposure to CP-506 (35.2 ± 23.7%) compared to vehicle exposure (18.8 ± 12.2%; P < 0.05). Furthermore, LNCaP *AR FANCA*^-/-^ tumors (P < 0.01), but not LNCaP *AR FANCD2*^-/-^ tumors (P = 0.16), exhibited significantly higher percentages of γH2AX positive cells when compared to LNCaP AR parental tumors in line with *in vitro* γH2AX results.

**Figure 4.**
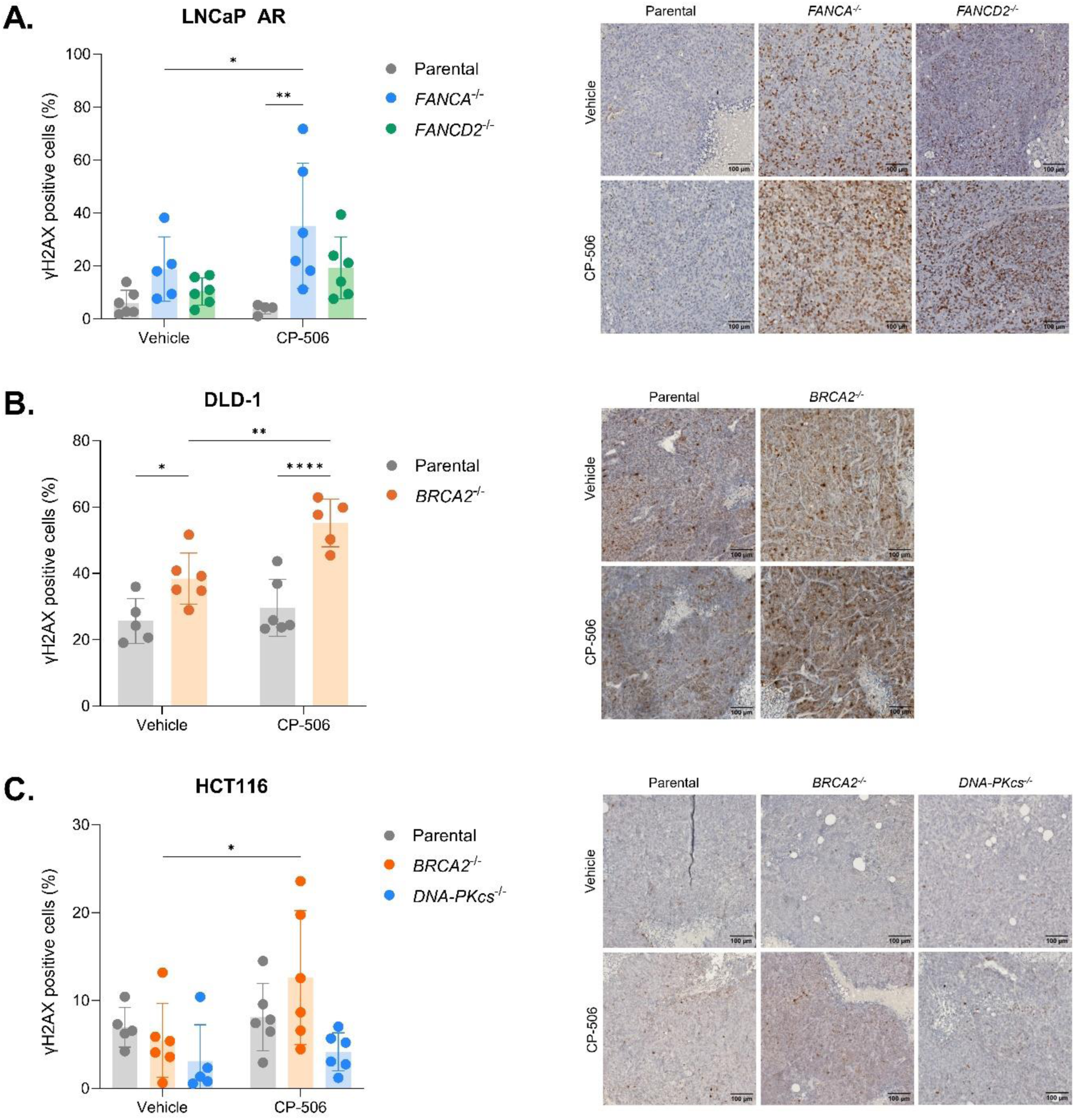
CP-506 induced persistent DSB damage in isogenic tumor xenografts deficient in FA or HR 48 hours post treatment. Percent of γH2AX positive cells in LNCaP AR (A), DLD-1 (B), and HCT116 (C) isogenic tumors with representative images of immunohistochemistry staining of γH2AX. n = 4-6 animals per group. Data are presented as mean ± SD. *: P < 0.05, **: P < 0.01, ****: P < 0.0001.

For DLD-1 tumors (Figure 4B), the baseline level of γH2AX positive cells was higher in BRCA2-deficient tumors (38.4 ± 7.7%) compared to parental tumors (25.6 ± 6.8%; P < 0.05). For parental tumors, no differences were observed in the amount of γH2AX positive cells after CP-506 exposure (29.7% ± 8.5%; P = 0.40) compared to vehicle-treated tumors. In contrast, the percentage of γH2AX positive cells in CP-506-treated DLD-1 *BRCA2^-/-^* tumors (55.2 ± 7.2%; P < 0.01) was significantly increased compared to vehicle exposure. Similar results were obtained for HCT116 tumors (Figure 4C). For HCT116 parental tumors, no significant difference was found between the vehicle-(7.0 ± 2.3%) and CP-506-treated (8.1 ± 3.8%; P = 0.67) tumors. For HCT116 *BRCA2^-/-^* tumors there was a significant increase in the percentage of γH2AX positive cells post CP-506 treatment (12.6 ± 7.6%) compared to the vehicle-treated HCT116 *BRCA2^-/^*^-^ tumors (5.5 ± 4.2%; P < 0.05), indicating residual DNA damage within these HR-deficient tumors. In line with the *in vitro* data (Supplementary Figure S7), HCT116 *DNA-PKcs^-/-^* tumors showed the lowest levels of γH2AX positive cells upon vehicle-(3.1 ± 4.2%) and CP-506-(4.2 ± 2.2%) treatment, with a slight, but not significant (P = 0.70) treatment-induced increase (Figure 4C). These results demonstrate that cancer cells and tumors deficient in FA or HR show a higher level of residual DNA damage 48 hours after CP-506 treatment as compared to their respective parental counterparts.

The alkaline comet assay confirms the presence of ICLs and DNA breaks upon CP-506 treatment To gain more insights into the type of CP-506-induced DNA damage and the potential underlying DNA repair mechanisms, the standard alkaline comet assay, to detect DNA strand breaks, and the modified alkaline comet assay, to detect ICLs, were performed. In the standard alkaline comet assay, the extent of DNA migration into the tail (%DNA in tail) is proportional to the number of DNA strand breaks, whereas in the modified alkaline comet assay, a decrease in %DNA in tail is indicative of an increase in ICL since these lesions inhibit DNA migration (Figure 5A) (23).

**Figure 5.**
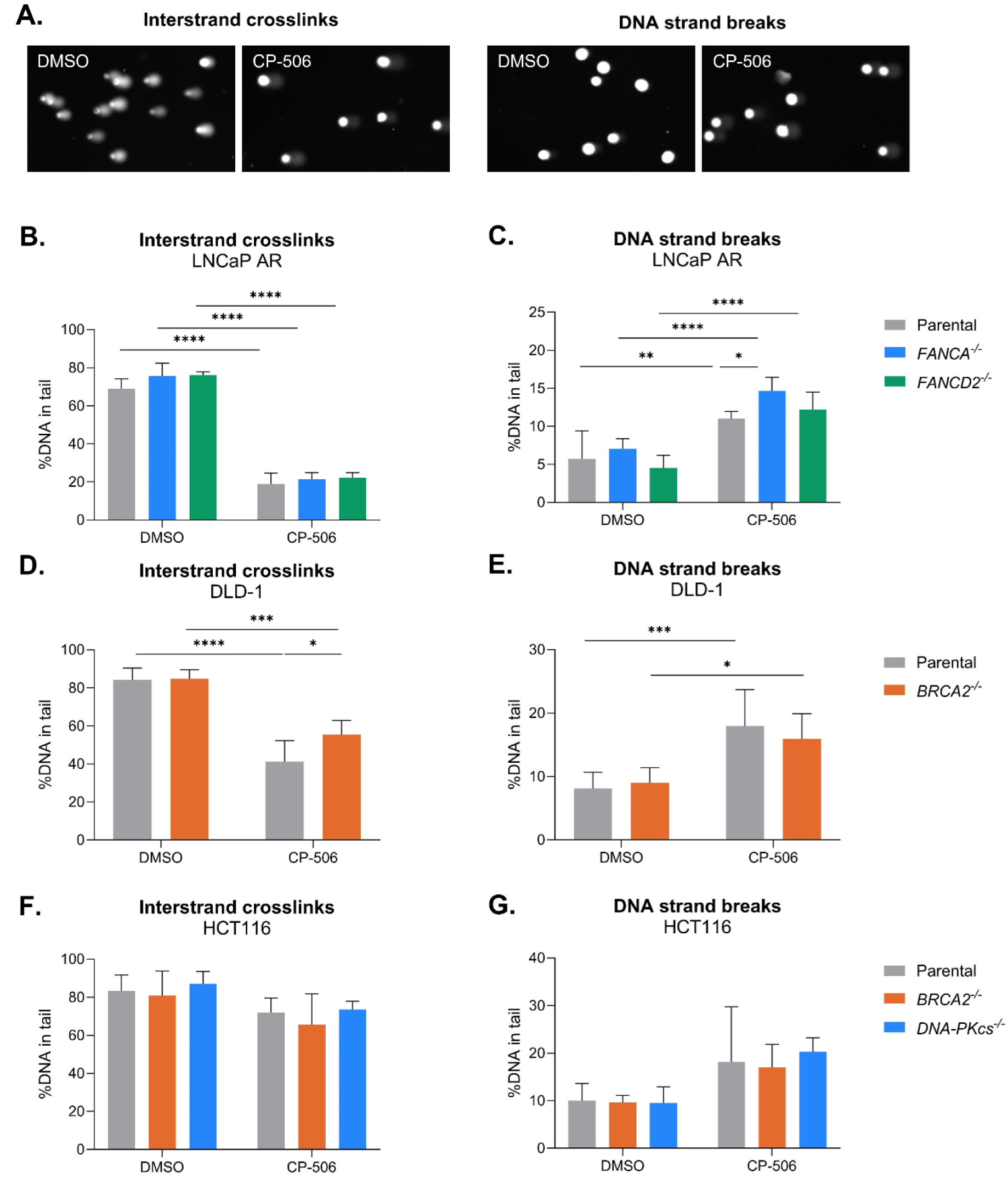
Anoxic exposure to CP-506 induced ICLs and DNA strand breaks in isogenic cancer cells 48 hours post treatment. Comet assay analysis of isogenic LNCaP AR, DLD-1, and HCT116 cells exposed to DMSO or CP-506 under anoxic conditions. Representative comets of cells exposed to CP-506 or DMSO (A). Cells were either assessed for interstrand crosslinks (ICLs; B, D, F) or DNA strand breaks (SSB and DSB; C, E, G). Medians from two biological repeats with one or more technical repeats were averaged ± SD. *: P < 0.05, **: P < 0.01, ***: P < 0.001, ****: P < 0.0001.

As expected, CP-506 induced ICLs, as shown by the reduction of the %DNA in tail compared to vehicle treatment for LNCaP AR (P < 0.0001), HCT116 (P = 0.27), and DLD-1 (P < 0.05) cancer cells at 48 (Figure 5B, 5D, 5F) and 72 (Supplementary Figure S8A, S8C, S8E) hours post start of treatment. At 48 and 72 hours post CP-506 treatment, DNA breaks were evident in all isogenic models (Figure 5C, 5E, 5G and Supplementary Figure S8B, S8D, S8F).

Upon CP-506 exposure of LNCaP AR cells, no differences in the reduction of the %DNA in tail, and thus presence of ICLs, were observed between FANCA-(21.3 ± 3.6%; P = 0.73) or FANCD2-deficient (22.2 ± 2.7%; P = 0.57) and parental cells (18.9 ± 5.7%; Figure 5B and Supplementary Figure S8A). DNA strand breaks were increased in FANCA-deficient cells (14.7 ± 1.8%; P < 0.05) compared to parental cells (11.0 ± 0.9 %DNA in tail; Figure 5C). This difference remained, although non-significant at 72 hours post treatment (Supplementary Figure S8B). In LNCaP AR *FANCD2^-/-^* cells, the amount of DNA strand breaks was not increased compared to parental cells upon CP-506 exposure (12.2 ± 2.3%; P = 0.69).

The presence of ICLs was significantly increased in DLD-1 parental cells as evidenced by the 14.2% (P < 0.05) and 16% (P < 0.01) reduction in %DNA in tail compared to BRCA2-deficient cells at 48 and 72 hours post CP-506 exposure, respectively (Figure 5D and Supplementary Figure S8C). DLD-1 *BRCA2^-/-^* cells displayed lower levels of DNA strand breaks as compared to DLD-1 parental cells at 48 (16 ± 4% vs 18 ± 5.7%; P = 0.38; Figure 5E) and 72 hours (20.4 ± 2.6% vs 25.3 ± 2.7%; P < 0.05; Supplementary Figure S8D). In HCT116 isogenic cancer cells, although showing a slight increase, the presence of ICLs or DNA strand breaks upon exposure to CP-506 was not significantly different compared to vehicle-treated cells (Figure 5F and 5G). There were no differences in the amount of CP-506-induced ICLs present in HCT116 parental (71.9 ± 7.6%) compared BRCA2-(65.6 ± 16.2%; P = 0.56) or DNA-PKcs-deficient (73.6 ± 65.6%; P = 0.96) cells (Figure 5F and Supplementary Figure S8E). There was no difference in the amount of DNA strand breaks in DNA-PKcs-deficient cells compared to parental cells at 48 hours (20.3 ± 2.9% vs 18.2 ± 11.5%; P = 0.87; Figure 5G) or 72 hours (20.5 ± 6% vs 12.5 ± 3.2%; P = 0.17; Supplementary Figure S8F) after exposure to CP-506. These findings confirm the formation of ICL and DNA damage upon CP-506 exposure, however, were unable to explain the differences in antitumor efficacy of CP-506 between isogenic xenograft models with deficiencies in various DNA repair pathways.

## Discussion

In the present study, we evaluated the role of DNA repair pathways in determining the cytotoxicity and antitumor effects of the hypoxia-activated DNA crosslinking agent CP-506. The FA and HR pathways are crucial for high-fidelity repair of ICLs and DSBs (9, 12). Here, we provided evidence that deficiencies in FA or HR, but not in NHEJ, repair pathways resulted in enhanced sensitivity to CP-506.

We first showed *in vitro* that FA- or HR-deficient cell lines were 2.0- to 3.4-fold more sensitive to CP-506 under anoxic conditions. Interestingly, HCT116 *DNA-PKcs^-/-^* cells deficient in NHEJ, a pathway also involved in repair of DSBs and associated with resistance to chemo- and radiotherapy (32, 33), were 1.6-fold more resistant to CP-506. Lastly, we confirmed in 3D *in vitro* hypoxic spheroid cultures, which more closely represents the tumor environment (34), that a deficiency in FA or HR, but not NHEJ, significantly increased the cytotoxicity of CP-506. Furthermore, we showed *in vivo* that CP-506 demonstrated monotherapeutic antitumor activity in all isogenic tumor xenograft models tested. In accordance with our *in vitro* data, CP-506 was more effective in delaying tumor growth in FA- and HR-deficient tumors compared to their parental tumors. Our findings are in line with previous studies demonstrating that cells or tumors deficient in FA or HR were more sensitive to HAPs such as PR-104 (19, 20) and TH-302 (20, 21) with similar mechanisms of action compared to CP-506. Furthermore, previous studies showed that the FA-deficient MDA-MB-468 triple-negative breast cancer xenograft model was highly sensitive to CP-506 with a 90% curative response (7). Lastly, FA- and HR-deficient tumors were also significantly more sensitive to chlorambucil (14, 35), a non-hypoxia-activated DNA crosslinking agent with a similar mechanism of action as CP-506, further evidencing ICL-induction to be the mechanism of action of CP-506. Altogether, these data further support the involvement of FA and HR in determining the cytotoxicity and antitumor effects of CP-506.

Mechanistically, we have demonstrated that CP-506-induced phosphorylation of H2AX was markedly increased in FA- or HR-deficient cancer cells and tumors compared to their respective parental counterparts. This is in agreement with other studies showing that in DNA repair proficient cells the levels of γH2AX foci restored to baseline level 48 hours after exposure to cisplatin or nitrogen mustard HN2, both non-hypoxia-activated ICL-inducing agents (36). Similarly, γH2AX foci remained persistent 48 hours post treatment in cancer cells deficient in HR (*XRCC3^-/-^*) or NER (*ERCC1^-/-^*) (36). Previous studies have observed that models with a high response to CP-506, including FaDu xenografts, showed persistent expression of γH2AX levels even 72 hours after exposure, while γH2AX levels reverted to baseline in models with a more resistant phenotype, including the UT-SCC-5 xenografts (29). Our results therefore suggest that parental cancer cells are able to repair the CP-506-induced DNA damage, contrary to FA- or HR-deficient cancer cells. Unlike radiation, the γH2AX response after exposure to ICL-inducing agents occurs with delayed kinetics, with a peak induction at 12-24 hours and gradual decrease until 48 hours (36-39). These delayed kinetics could be explained by need of cells to progress into the S-phase before γH2AX induction (11) or by other causes, i.e. not related to DNA repair.

Phosphorylation of H2AX on Ser139 is widely known as a marker for DSB, however, its functionality extends beyond its role in DNA repair. While H2AX is phosphorylated at sites of DSBs, its presence does not necessarily indicate the occurrence of DSBs (40). Increased or persistent γH2AX expression has been found in situations of replication stress due to replication fork stalling or damage (41-43), apoptosis (44-47), or cell senescence (48). To unravel the underlying cause of the persistent γH2AX expression in FA- and HR-deficient cells upon CP-506 treatment, we performed the alkaline comet assays to assess the presence of ICLs and DNA strand breaks. Considering the essential role of the FA pathway in the sensing and coordinating the unhooking of ICLs (49, 50), defects within the FA pathway were expected to result in the accumulation of ICLs and hence less ICL-induced DSBs and associated phosphorylation of H2AX. Contrary to this hypothesis, we observed persistent γH2AX expression in FA-deficient cells *in vitro* and *ex vivo*. The enhanced phosphorylation in FA-deficient cells could be caused by stalled replication forks due to the occurrence of ICLs (43). The presence of γH2AX at stalled replication forks is required for the recruitment and accumulation of FANCD2, playing a crucial role in promoting replication fork restart (51, 52). However, we did not observe differences in the presence of ICLs between LNCaP AR parental and FA-deficient cells. Besides, the persistent γH2AX levels observed in FA-deficient cells could also be attributed to the formation of passive DSBs at the site of a stalled DNA polymerase upon the encounter of an ICL (53). This is supported by our findings, showing an increase in DNA strand breaks as detected by the alkaline comet assay in FA-deficient cells compared to their parental cells after CP-506 treatment. Despite the vast evidence of the involvement of the FA pathway in the sensing and unhooking of ICLs, our results are in line with previous studies suggesting that the FA pathway is to a greater extent involved in the downstream DSB repair process (54, 55). Moreover, it has been suggested that the NER pathway is responsible for the detection and incision of ICLs formed upon exposure to crosslinking agents (10, 55-57), indicating that the removal of ICL is not solely dependent on the FA pathway. Therefore, more research is needed to confirm the role of the NER pathway in the removal of ICLs induced by CP-506.

The modified alkaline comet assay confirmed the presence of CP-506-induced ICLs in all isogenic cell lines, in line with our previously published *in vitro* and *in vivo* work showing that CP-506 caused induction of ICLs and DSBs specifically under severe hypoxic conditions (7, 8). Interestingly, we observed that the amount of ICLs in CP-506-treated cells did not revert to vehicle levels after 72 hours. Therefore, the lack in observed differences in ICL repair between isogenic models, especially the LNCaP AR model, could reflect a delay in the repair kinetics of CP-506 induced-ICLs, which may require more than 72 hours to resolve. This is supported by previous studies, which provided evidence for distinct unhooking and repair mechanisms of ICLs induced by different crosslinking agents (58). In case of BRCA2-deficient cells, we hypothesized delayed repair of ICL-repair mediated DSB-intermediates upon CP-506 exposure. Contrary to what we hypothesized, we did not observe any differences in the presence of DNA strand breaks in HCT116 cells, whereas a decrease was observed in DLD-1 BRCA2^-/-^ cells compared to their parental cells. Unexpectedly, DLD-1 parental cells also contained more ICLs compared to BRCA2-deficient cells. A possible explanation could be that due to the genetic inhibition of BRCA2 other genetic pathways are upregulated, resulting in better DNA repair compared to parental cells. This difference, however, was not observed for HCT116 cells. Interestingly, the more resistant NHEJ-deficient model showed at both time points the highest level of DNA strand breaks compared to HR-deficient and parental HCT116 cells. It is important to note that the alkaline comet assay is unable to differentiate between SSBs or DSBs (59). Furthermore, the effector metabolites of CP-506 are bifunctional alkylating agents, also inducing monoadducts and intrastrand crosslinks in addition to ICLs and ICL-repair DSB-intermediates (7, 8), which might complicate the interpretation of our results. One should note that high concentrations of CP-506 could lead to cytotoxic effects, which may have introduced an underestimation in the detection of ICLs and DNA damage using the alkaline comet assay, although lower concentrations did not produce detectable levels of ICLs. Overall, our data support γH2AX expression as a more sensitive marker of treatment response to cytotoxic concentrations of CP-506, as opposed to the alkaline comet assays, which showed variability and were unable to provide a more mechanistic understanding on the differences in therapeutic sensitivity between isogenic cell lines. Future studies should, therefore, include additional markers of apoptosis, necrosis, and cell cycle to discriminate between persistent γH2AX due to the inability of cells to repair DSB or collapsed replication forks or whether the persistent phosphorylation of H2AX is associated with different cellular processes. Additionally, use of other markers of DNA damage such as RAD51 and 53BP1, which are directly related to HR (60-63), and incorporating DNA adductomics analyses (64) to directly measure DNA adducts could further increase our understanding of kinetics of DNA damage induction and repair after CP-506 exposure.

As previously shown by published studies (7), tumor hypoxia is an important but nor sole determinant of CP-506 antitumor efficacy. While tumor hypoxia was confirmed in the isogenic models in the present study, those exhibiting the greatest antitumor response to CP-506 did not display the largest HF, suggesting that tumor hypoxia alone does not fully account for the differential responses to CP-506 treatment. These findings are consistent with the hypothesis that the intrinsic sensitivity – i.e., the HRD status – of tumor cells plays a major role in determining the ultimate efficacy of CP-506, given that the tumors are hypoxic (5). In line with the enhanced sensitivity of HRD cells to ICL-inducing chemotherapies (14-16), PARPi (17, 18), PR-104 (19, 20), and TH-302 (20, 21), our results suggests that CP-506 similarly leverages synthetic lethality as a therapeutic strategy. This highlights the importance of HRD status as a clinical biomarker for patient stratification. Pan-cancer analyses have shown that HRD is prevalent not only in ovarian and breast cancers, but also in other malignancies such as for example prostate, pancreatic, and endometrial cancer. However, reported frequencies varied widely (6-20%) (65-68), which can be attributed to methodology – such as whether HRD was defined by mutational status of HR-related genes or genomic scars as reviewed in (69) – and biological variability including inter-patient differences within the same cancer type (68) or the presence of reversion mutations within individual patients (70). This underlines the importance of further exploring genetic HRD testing to improve patient stratification and guide treatment decisions in future clinical applications.

Despite the promising results of CP-506 as monotherapy, its therapeutic efficacy is expected to be the greatest when combined with complementary treatment modalities targeting well-oxygenated tumor cells (71), similarly as has been proposed for other HAPs, including TH-302 (20, 72-75), PR-104 (20, 76), and tirapazamine (77-80). Moreover, CP-506 has been proven to enhance the therapeutic efficacy of radiation, particularly using hypofractionation schedules that do not allow for reoxygenation (29). However, these published studies showed variability in the therapeutic response of combining radiation with CP-506 between different tumor models. This is likely attributable to differences in the intrinsic sensitivity of the models, as suggested by our data and evidenced by the differential response to CP-506 and mitomycin c in cell viability assays (29). Given the role of hypoxia in immunotherapy resistance (81-83), targeting tumor hypoxia with CP-506 may help to overcome this challenge, enhancing immunotherapy treatment efficacy, as previously has been shown for TH-302 (82, 84, 85). Moreover, exploring this combination in HRD tumors would be of interest since these tumors show high genomic instability and mutational burden (86-88), resulting in an increased neoantigen load (89). Clinical studies have shown that HRD tumors may respond better to PD-1/PD-L1 and/or CTLA-4 immune checkpoint inhibitors (90, 91). Further studies are warranted to investigate the potential of CP-506 in combination with immunotherapy in these contexts.

In conclusion, CP-506 is a novel hypoxia-activated DNA crosslinking agent that is selectively activated only under severe hypoxic conditions. Several HAPs have previously been evaluated in both preclinical and clinical settings, but despite promising preclinical findings, implementation of HAPs into the clinic has yet to be successful, with a lack of patient stratification at least in part accountable for this failure. Identification of key factors influencing the tumoral response to HAPs is therefore essential for their successful clinical application. Here, we demonstrated in several *in vitro* and *in vivo* models that isogenic cells and tumors deficient in FA or HR, but not in NHEJ, are markedly more sensitive to CP-506. Based on these findings, we propose that CP-506 is expected to be most effective in tumors that are both hypoxic and FA- and/or HR-deficient. CP-506 is currently being evaluated in an ongoing phase I/IIa clinical trial (NCT04954599), in which tumor hypoxia and DNA repair status will be assessed.

## Data availability

The datasets generated and analyzed during the current study are available from the corresponding author on reasonable request. A publicly available implementation of the DynUNet architecture was used to support an initial step in image preprocessing (https://docs.monai.io/en/0.7.0/_modules/monai/networks/nets/dynunet.html). This architecture was adapted using a training script developed in-house, which can be shared upon reasonable request for academic purposes.

## Acknowledgements

We acknowledge Jeff Smaill and Adam Patterson as inventors of CP-506 and thank them for their contributions to its development (25). Furthermore, we would like to acknowledge the technical support of Hellen Steinbusch and Prof. Mario Losen from the Department of Psychiatry and Neuropsychology, School for Mental Health and Neuroscience, Maastricht University and Laura Peeters from the Department of Orthopedic Surgery, Maastricht University with microscope image acquisition and slide scanning. This work was financially supported by the ERC Advanced Grant HYPOXIMMUNO (ERC-ADG-2015, no. 694812), the ERC Proof of Concept grant “Reverse the Advantage” (ERC-2022-PoC2-101082238), and two awarded travel grants: the Klaas Breur Travel Award granted by the Netherlands Society for Radiobiology (NVRB) and a travel grant awarded by the European Association for Cancer Research (EACR).

## Conflict of interest statement

PL reports, within and outside of the scope of the current manuscript, grants or sponsored research agreements from Radiomics SA, Concert Pharmaceuticals SA, and LivingMed Biotech srl. He received a fee and/or reimbursements (in cash or in kind) for presentations, consultancy, or travel from AstraZeneca, BHV srl, and Roche. PL currently holds or has held minority shares in Radiomics SA, Convert Pharmaceuticals SA, Comunicare SA, LivingMed Biotech srl, and Bactam srl. PL is listed as a co-inventor on several patents: two issued patents with royalties on radiomics (PCT/NL2014/050248 and PCT/NL2014/050728), licensed to Radiomics SA; one issued patent on mtDNA (PCT/EP2014/059089), licensed to ptTheragnostic/DNAmito; one issued patent on LSRT (PCT/P126537PC00, US Patent No. 12,102,842), licensed to Varian; one issued patent on a radiomic hypoxia signature (U.S. Patent 11,972,867), licensed to a commercial entity; one issued Prodrug-related patent (WO2019EP64112) without royalties; one pending, unlicensed patent on Deep Learning-Radiomics (N2024889); and three non-patented software inventions, licensed to ptTheragnostic/DNAmito, Radiomics SA, and Health Innovation Ventures. PL declares that none of these entities had any involvement in the preparation of this manuscript. LJD holds, within the submitted work, minority shares in the companies Convert Pharmaceuticals SA, and, outside of the submitted work, minority shares in LivingMed Biotech srl. He is also co-inventor on a granted patent on LSRT (PCT/P126537PC00, US Patent No. 12,102,842), licensed to Varian. Similarly, JT has minority shares in Convert Pharmaceuticals SA. The authors confirm that none of the entities listed above were involved in the preparation of this manuscript. All other authors report no conflict of interest

## Author contributions

**L. Schuitmaker:** Data curation, formal analysis, investigation, methodology, software, validation, visualization, writing – review and editing. **A.M.A. van der Wiel:** Data curation, formal analysis, investigation, methodology, validation, visualization, writing – original draft, writing – review and editing. **N.G. Lieuwes:** Formal analysis, investigation, methodology. **R. Biemans:** Formal analysis, investigation. **N.A.M. Mutsters:** Formal analysis, investigation, validation. **J. Jung:** Formal analysis, investigation. **V. Claudino Bastos:** Data curation, investigation, methodology, validation. **E. Prades Sagarra:** Investigation, writing – review and editing. **S. Kuang**: Methodology, software, writing – review and editing. **S.A.S. Langie**: Methodology, resources, validation, writing – review and editing. **J. Setton**: Resources. **J. Theys**: Conceptualization, supervision, writing – review and editing. **A. Yaromina:** Conceptualization, formal analysis, investigation, methodology, supervision, software, validation, writing – original draft, writing – review and editing. **L.J. Dubois:** Conceptualization, funding acquisition, investigation, methodology, project administration, resources, supervision, validation, visualization, writing – original draft, writing – review and editing. **P. Lambin:** Conceptualization, funding acquisition, project administration, resources, supervision, validation, visualization, writing – original draft, writing – review and editing.

