## Supplementary information for "Deficiencies in the Fanconi Anemia or the Homologous Recombination pathway enhance the antitumor effects of the novel hypoxia-activated prodrug CP-506"

### Supplementary Materials and Methods

#### Vital tumor segmentation using Deep-learning DynUNet models

A DynUNet (1) architecture (implemented using Python 3.11.5 and Monai 1.3.0) was employed to delineate vital tumor areas, excluding necrotic tumor areas, connective tissues, and processing or staining artefacts, within the scanned slides. For the input to the DynUNet architecture, two observers (N.G.L. and L.S.) manually annotated a randomly selected subset of 30 scanned slides in RGB format using ImageJ software 1.54f.

The dataset was stratified into a training set consisting of 22 images and a validation set comprising 8 images ensuring equal contribution of the different isogenic tumor models. To reduce the computational complexity, the RGB images were first down sampled by a factor of 5 using bilinear interpolation. During training, 32 patches (512 x 512 pixels) were randomly extracted from each image and data augmentation techniques, including random adjustments in contrast, intensity, rotation, and flip, were applied.

For training the DynUNet, a deep supervision mechanism was incorporated, ensuring robust learning at multiple network depths. A combined Dice and cross-entropy loss was employed to optimize segmentation performance. The Adam optimizer was used for 1000 epochs with an initial learning rate of 0.002, following a polynomial decay schedule. The model with the best validation performance was retrained for final interference.

During inference, a sliding window approach with a stride of 384 was used to ensure comprehensive coverage and precision in the prediction of vital areas. The outputs from overlapping patches were integrated using Gaussian fusion. The final segmentation mask was obtained by applying a threshold of 0.5 to the prediction probabilities, after which the mask was up sampled back to match the original image resolution using nearest interpolation.

#### Assessment of tumor hypoxia by pimonidazole immunofluorescence

To assess the hypoxic fraction (HF), tumor sections were stained for the hypoxia marker pimonidazole (NPI, Inc) as previously described (6, 22). In short, frozen tumor sections (7 µm) were fixed in cold acetone and blocked with 5% (v/v) normal goat serum in antibody diluent. Overnight incubation (4 °C) with rabbit anti-pimonidazole antibody (1:250 in antibody diluent; NPI, Inc.) was followed by TBS-Tw washes and a one-hour incubation with goat anti-rabbit IgG Alexa Fluor 488 (1:500 in antibody diluent; Invitrogen) at RT. Sections were incubated with DAPI for 10 minutes and mounted in fluorescent mounting medium (DakoCytomation). Images were acquired with an Olympus BX51WI microscope equipped with a 10x objective, Hamamatsu EM-CCD C9100 digital camera, and a Ludl Mac 2000 motorized stage. For quantitative image analysis, viable tumor tissue was delineated manually based on DAPI with ImageJ (N.G.L. and R.B.) blinded to the subject coding. Thereafter, thresholds were set manually to distinguish pimonidazole fluorescence from background and HF was calculated as the ratio of the pimonidazole positive area to the viable tumor area.

### Supplementary Figures

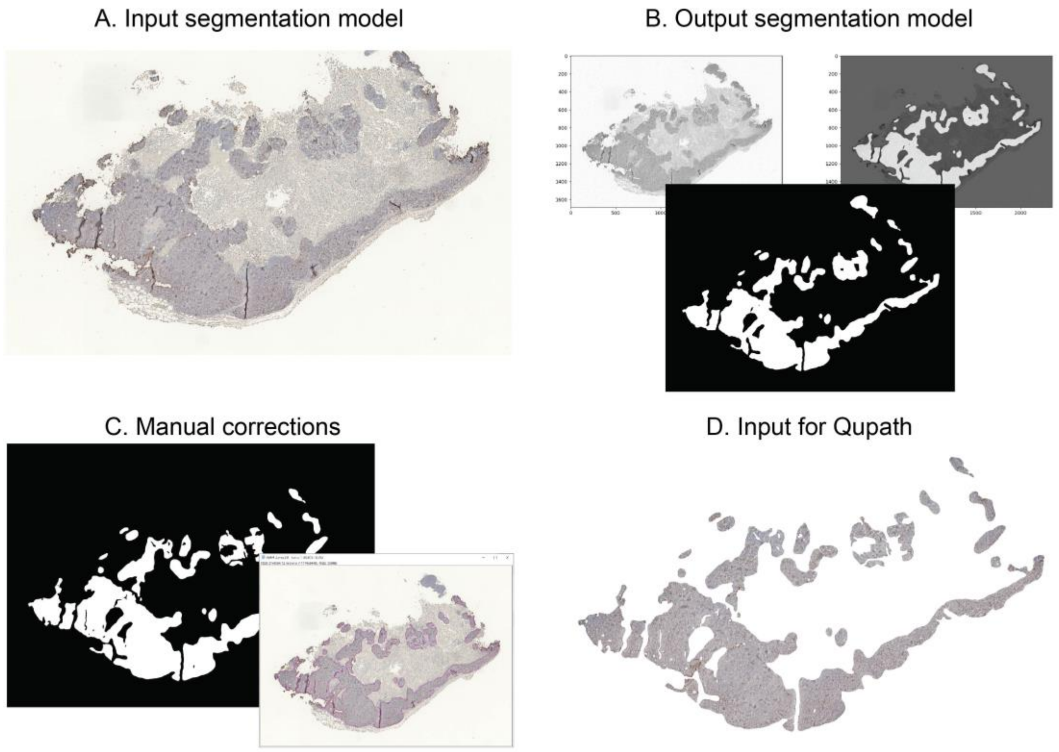


**Supplementary Figure S1.** Workflow for the segmentation of vital masks of isogenic tumor sections for the analysis of γH2AX immunohistochemistry staining. A deep-learning DynUNet model was generated to segment the vital tumor areas from raw RGB images. After performing manual corrections of the segmentations using ImageJ version 1.54f, the corrected vital mask was overlaid with the raw RGB image, which generates an input image in Qupath version 0.4.3. for immunohistochemistry analysis of γH2AX (% positive cells).

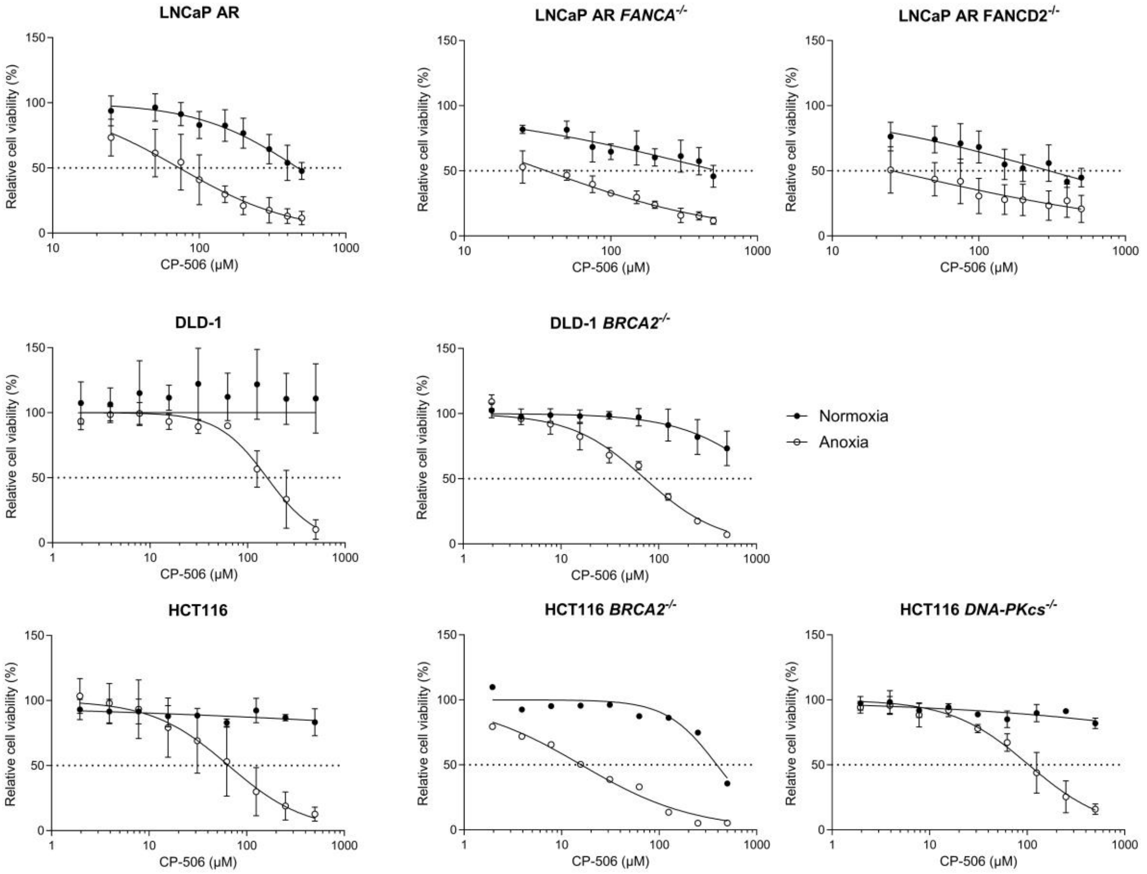


**Supplementary Figure S2.** Cell viability dose-response curves upon CP-506 exposure in isogenic cancer cell lines proficient or deficient in the FA *(FANCA$^{-/-}$* or *FANCD2$^{-/-}$)*, HR *(BRCA2$^{-/-}$)*, or NHEJ *(DNA-PKcs$^{-/-}$)* DNA repair pathways. Isogenic cancer cells were exposed to increasing CP-506 concentrations under normoxic (closed circles) and anoxic (open circles) conditions after which cell viability was assessed and $IC_{50}$ values were determined.

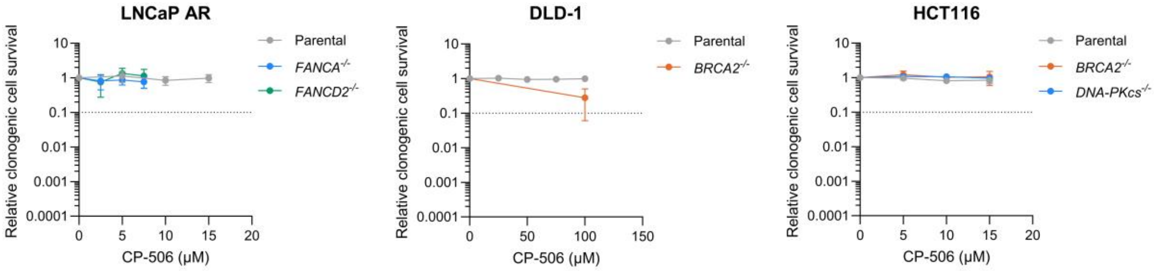


**Supplementary Figure S3.** Clonogenic cell survival after 4 hours of exposure to CP-506 of isogenic cancer cell lines proficient or deficient in DNA repair pathways upon normoxic conditions.

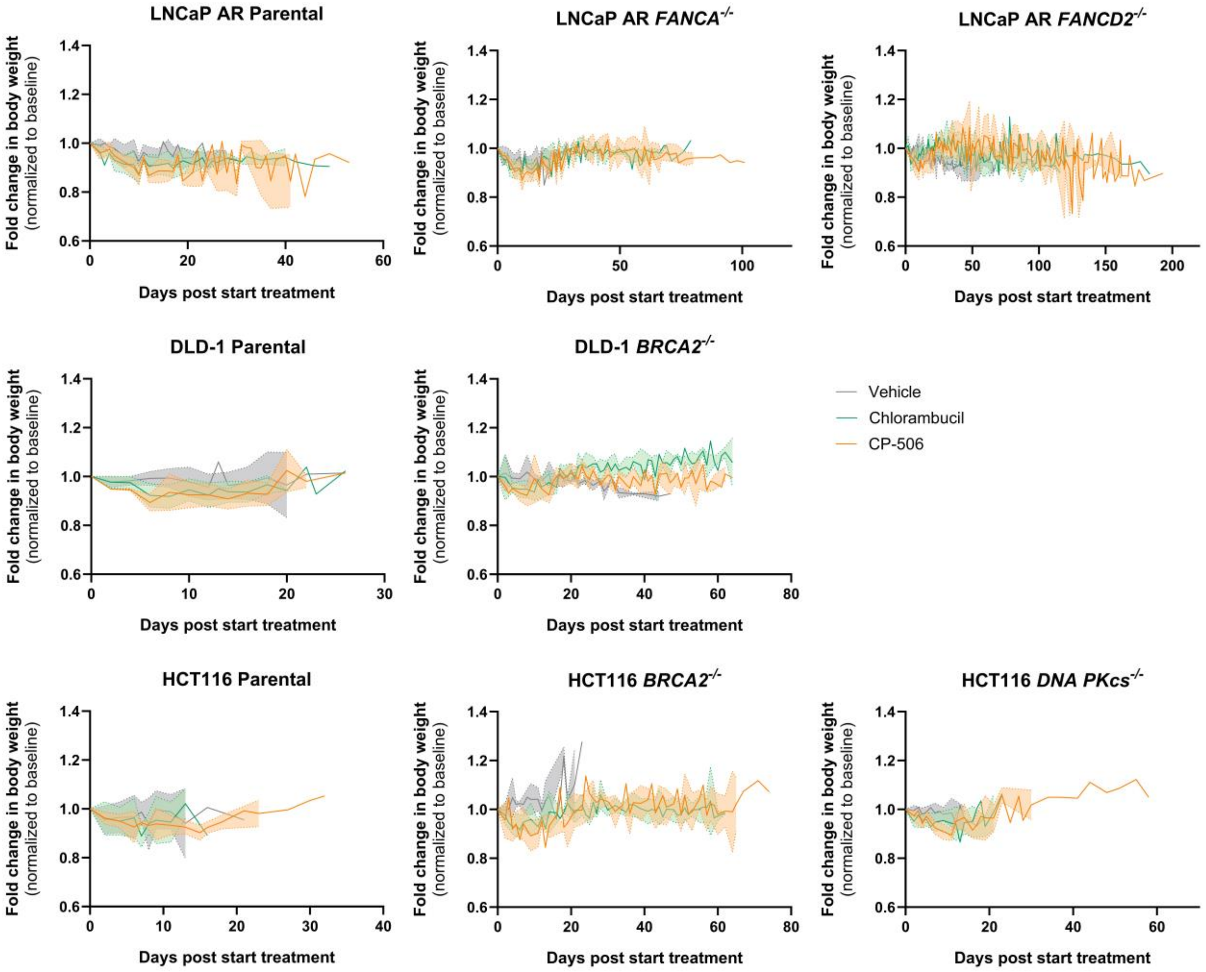


**Supplementary Figure S4**. Effect of vehicle, chlorambucil, or CP-506 treatment on the body weight of mice bearing isogenic tumor xenografts. Body weight changes over time upon vehicle (WFI), chlorambucil, or CP-506 treatment, normalized to body weight at the first day of treatment. Data are shown as mean ± SD (n = 8-10 animals per group).

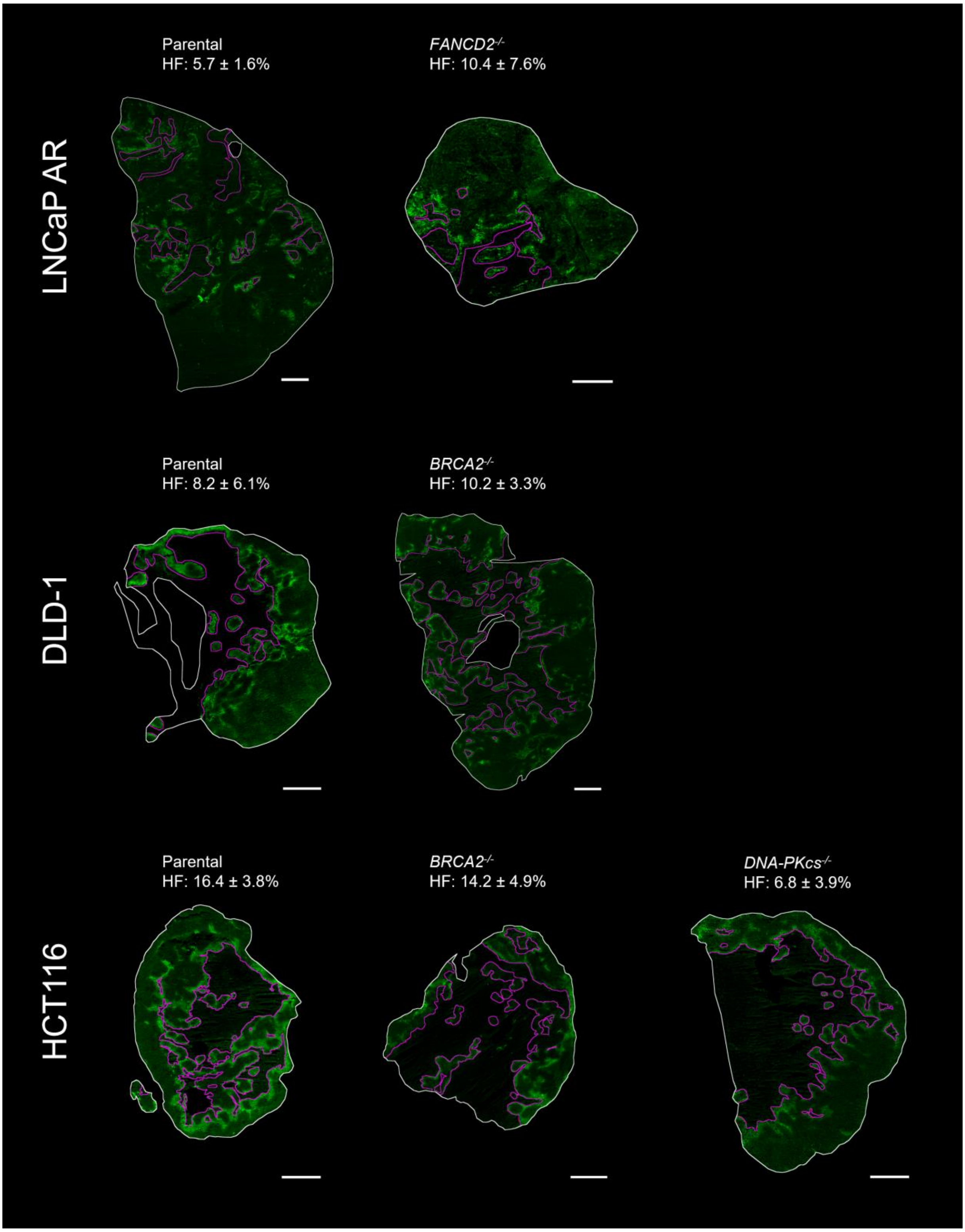


**Supplementary Figure S5**. Pimonidazole hypoxic fraction (HF) in isogenic xenografts 6 hours post vehicle treatment. Representative images showing total tumor area (white) and necrotic tumor area (magenta); background outside the total tumor area was cleared for visualization. Data are reported as mean ± SD (n = 4-6 animals per group). Scale bar represents 1 mm.

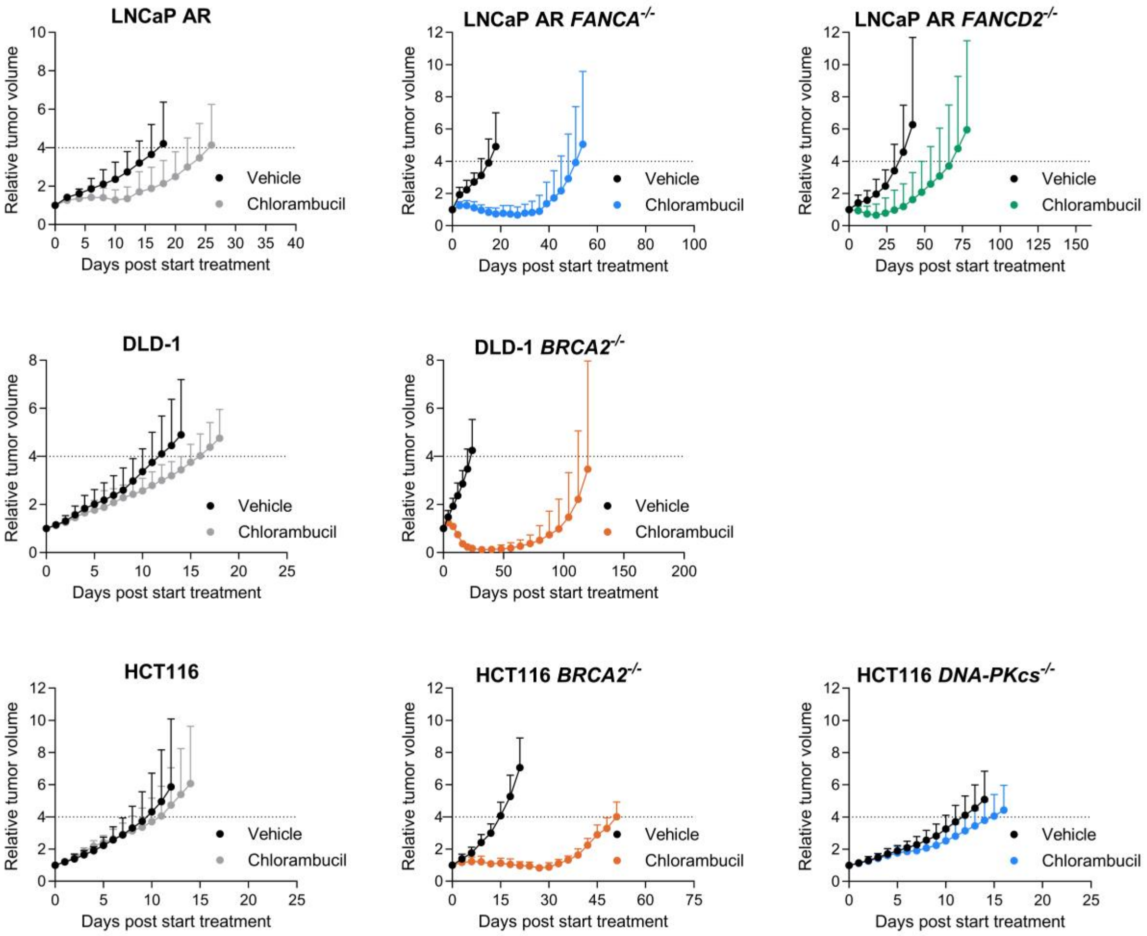


**Supplementary Figure S6.** Antitumor effects of vehicle (WFI) or chlorambucil treatment (3 mg/kg, QD5, IP) in mice bearing isogenic tumor xenografts, proficient or deficient in FA (*FANCA*$^{-/-}$ or *FANCD2*$^{-/-}$), HR (*BRCA2*$^{-/-}$), or NHEJ (*DNA-PKcs*$^{-/-}$). Data are presented as mean ± SD (n = 8-10 animals per group).

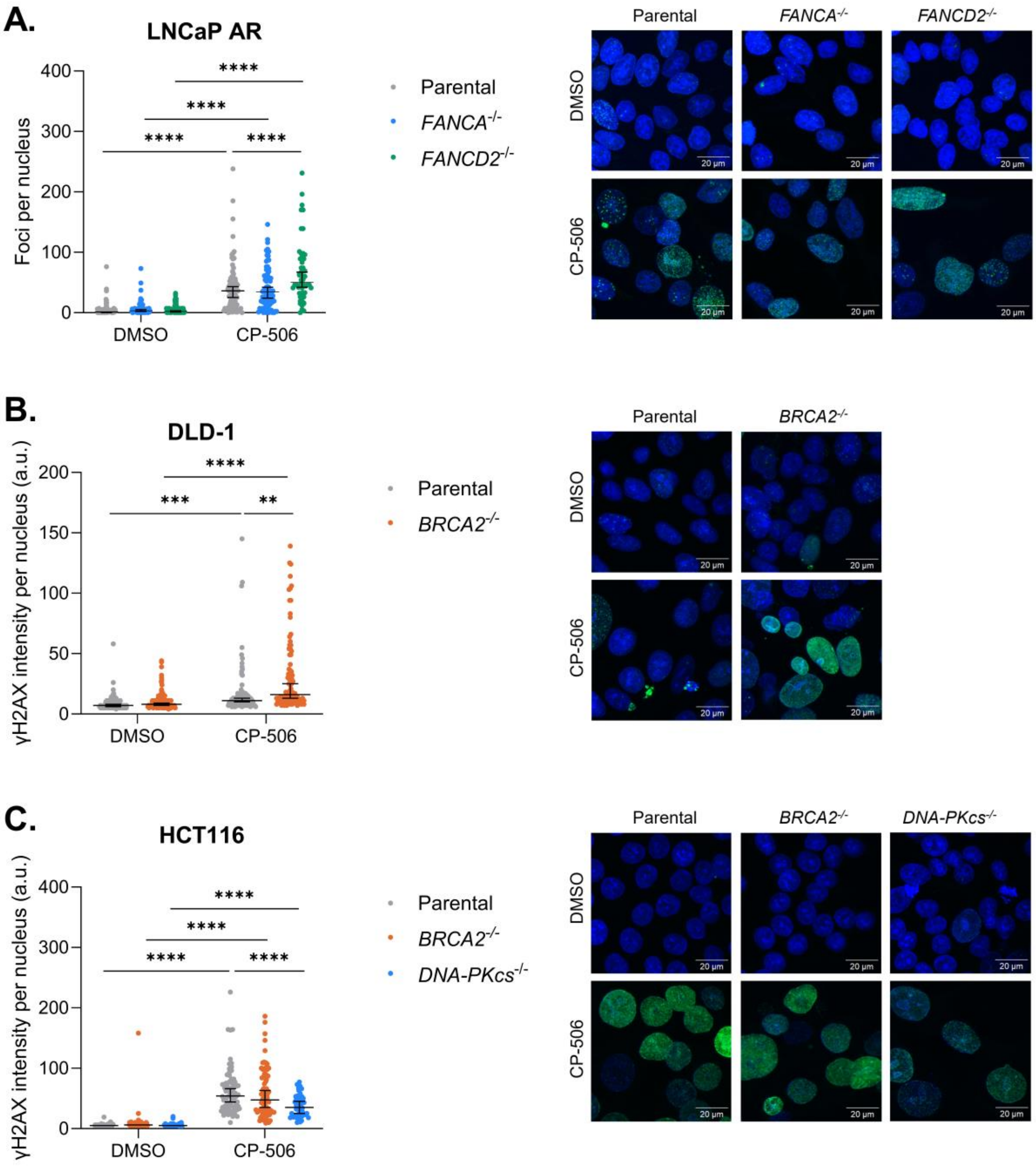


**Supplementary Figure S7**. CP-506 induced persistent DNA damage in FA- and HR-deficient isogenic cancer cells under anoxic conditions as determined by immunofluorescence 72 hours post start of treatment. Foci count per nucleus for LNCaP AR isogenic cancer cells (A) and quantification of γH2AX immunofluorescence intensity per nucleus for DLD-1 (B) and HCT116 (C) isogenic cancer cells with representative γH2AX immunofluorescence images. Blue: Hoechst; green: γH2AX. n ≥ 57 cells per condition and data are presented as median (IQR). *: P < 0.05, **: P < 0.01, ***: P < 0.001, ****: P < 0.0001.

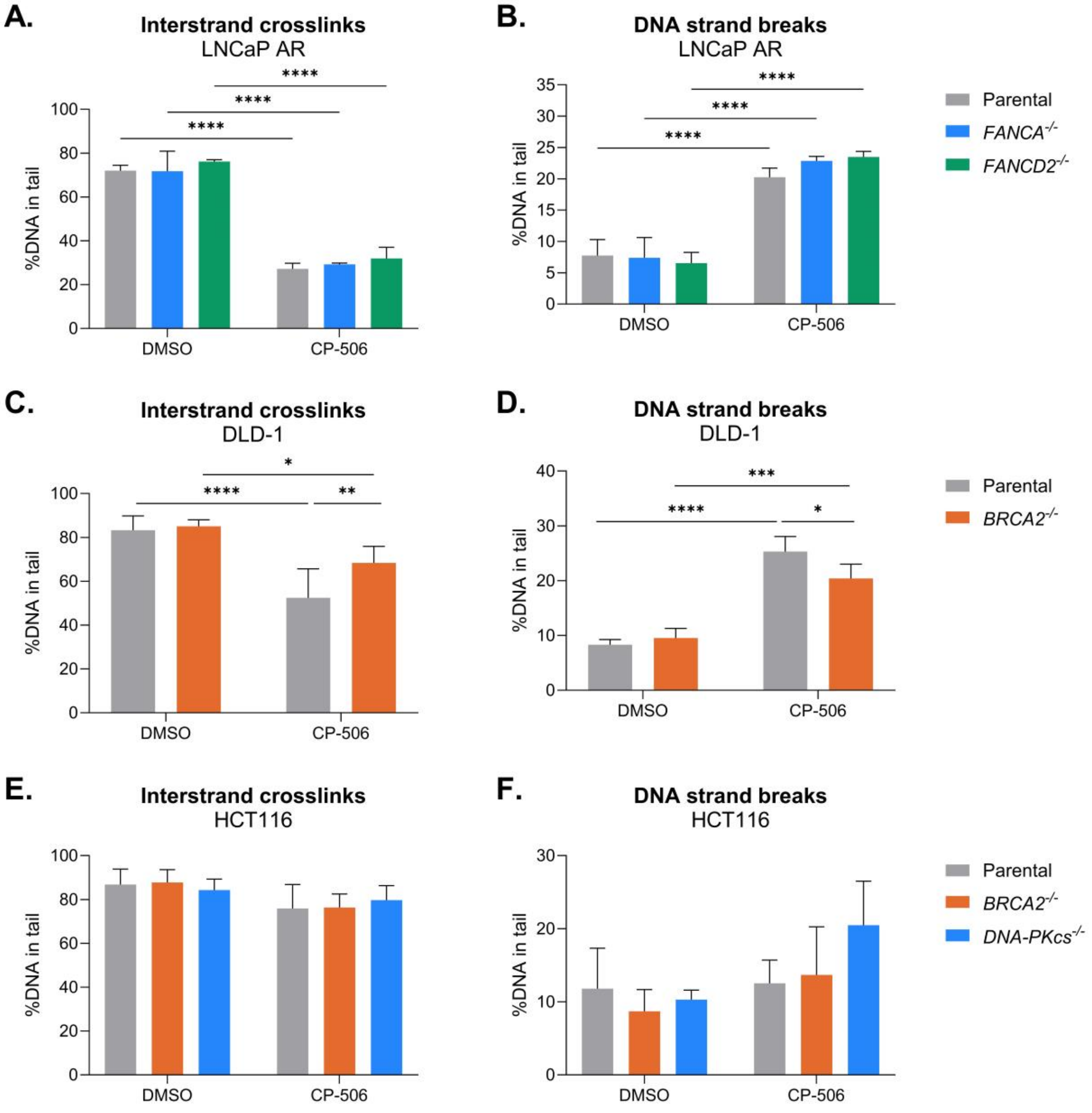


**Supplementary Figure S8.** Anoxic exposure to CP-506 induced ICLs and DNA strand breaks in isogenic cancer cells 72 hours post treatment. Comet assay analysis of isogenic LNCaP AR, DLD-1, and HCT116 cells exposed to DMSO or CP-506 under anoxic conditions. Cells were either assessed for interstrand crosslinks (ICLs; A, C, E) or DNA strand breaks (SSBs and DSBs; B, D, F). Medians from two biological repeats with one or more technical repeats were averaged ± SD. *: $P < 0.05$, **: $P < 0.01$, ***: $P < 0.001$, ****: $P < 0.0001$.

### Supplementary Tables

**Supplementary Table S1.**

An overview of all isogenic cell lines, their tissue of origin, genetic mutation and corresponding DNA repair pathway affected, provider, and culture medium used.

| **Cell line** | **Cancer type** | **DNA repair pathway** | **Provider** | **Culture medium** |
|---|---|---|---|---|
| LNCaP AR | Prostate | Parental | MSKCC | DMEM, supplemented with 10% FBS |
| LNCaP AR *FANCA*$^{-/-}$ | Prostate | FA | MSKCC | DMEM, supplemented with 10% FBS |
| LNCaP AR *FANCD2*$^{-/-}$ | Prostate | FA | MSKCC | DMEM, supplemented with 10% FBS |
| DLD-1 | Colorectal | Parental | Horizon Discovery | McCoy's 5A, supplemented with 10% FBS |
| DLD-1 *BRCA2*$^{-/-}$ | Colorectal | HR | Horizon Discovery | McCoy's 5A, supplemented with 10% FBS |
| HCT116 | Colorectal | Parental | Horizon Discovery | DMEM, supplemented with 10% FBS |
| HCT116 *BRCA2*$^{-/-}$ | Colorectal | HR | Ximbio | DMEM, supplemented with 10% FBS |
| HCT116 *DNA-PKcs*$^{-/-}$ | Colorectal | NHEJ | Horizon Discovery | DMEM, supplemented with 10% FBS |

FA: Fanconi anemia pathway; HR: homologous recombination; NHEJ: non-homologous end joining; FANCA: FA complementation group A; FANCD2: FA complementation group D2; BRCA2: breast cancer type 2 susceptibility protein; DNA-PKcs: DNA-dependent protein kinase, subunit C; MSKK: Memorial Sloan Kettering Cancer Center; DMEM: Dulbecco's Modified Eagle Medium; FBS: fetal bovine serum.

**Supplementary table S2**

Read-out parameters monotherapeutic efficacy CP-506 in isogenic xenograft models.

| Cell line | Cancer type | DNA repair pathway | Mouse strain | T4xSV – vehicle | T4xSV – CP-506 | ER | TGI | P-value T4xSV vehicle vs CP-506 | P-value ER vs WT | P-value TGI vehicle vs CP-506 |
|---|---|---|---|---|---|---|---|---|---|---|
| LNCaP AR | Prostate | Parental | NOD.Cg-Prkdc$^{SCID}$ Il2rg$^{tm1Wjl}$/SzJ | 19.2 ± 5.8 | 29.4 ± 9.0 | 1.5 ± 0.5 | 52.3 ± 56.4% | 0.29 | - | < 0.05 |
| LNCaP AR *FANCA*$^{-/-}$ | Prostate | FA | | 16.6 ± 4.2 | 66.7 ± 18.3 | 4.0 ± 1.1 | 98.6 ± 1.5% | < 0.0001 | < 0.0001 | < 0.0001 |
| LNCaP AR *FANCD2*$^{-/-}$ | Prostate | FA | | 40.6 ± 13.4 | 139.0 ± 30.5 | 3.4 ± 0.8 | 97.7 ± 3.2% | < 0.0001 | < 0.0001 | < 0.0001 |
| DLD-1 | Colorectal | Parental | BALB/c nu/nu | 13.9 ± 4.6 | 18.3 ± 2.8 | 1.3 ± 0.2 | 37.9 ± 10.3% | 0.49 | - | < 0.01 |
| DLD-1 *BRCA2*$^{-/-}$ | Colorectal | HR | | 25.5 ± 7.8 | 73.5 ± 17.9 | 2.9 ± 0.7 | 77.8 ± 11.0% | < 0.0001 | < 0.0001 | <0.0001 |
| HCT116 | Colorectal | Parental | NU-Foxn1 nu/nu | 10.2 ± 3.8 | 17.0 ± 5.7 | 1.7 ± 0.6 | 59.5 ± 12.9% | < 0.01 | - | < 0.01 |
| HCT116 *BRCA2*$^{-/-}$ | Colorectal | HR | | 15.3 ± 2.3 | 61.4 ± 9.2 | 4.0 ± 0.6 | 95.7 ± 3.2% | < 0.0001 | < 0.0001 | < 0.0001 |
| HCT116 *DNA-PKcs*$^{-/-}$ | Colorectal | NHEJ | | 12.6 ± 2.7 | 17.5 ± 3.6 | 1.4 ± 0.3 | 33.2 ± 17.3% | < 0.05 | 0.18 | 0.15 |

T4xSV: time to reach 4 times starting volume; ER: enhancement ratio; TGI: tumor growth inhibition at the day which respective vehicle-treated tumors reached four times starting volume (4xSV).

**Supplementary Table S3**

An overview of the optimized settings per isogenic tumor model for positive cell detection in QuPath version 0.4.3

| Parameter | LNCaP AR | DLD-1 | HCT116 |
|---|---|---|---|
| **Setup parameters** | | | |
| Detection image | Optical density sum | Optical density sum | Optical density sum |
| Requested pixel size | 0.5 µm | 0.5 µm | 0.5 µm |
| **Nucleus parameters** | | | |
| Background radius | 85.0 µm | 8.0 µm | 50.0 µm |
| Use opening by reconstruction | TRUE | FALSE | FALSE |
| Medium filter radius | 0.0 µm | 0.0 µm | 0.0 µm |
| Sigma | 2.0 µm | 1.4 µm | 1.3 µm |
| Minimum area | 20.0 $\mu m^2$ | 10.0 $\mu m^2$ | 13.0 $\mu m^2$ |
| Maximum area | 400.0 $\mu m^2$ | 400.0 $\mu m^2$ | 400.0 $\mu m^2$ |
| **Intensity parameters** | | | |
| Threshold | 0.26 | 0.16 | 0.16 |
| Max background intensity | 10.0 | 2.0 | 2.0 |
| Split by shape | TRUE | TRUE | TRUE |
| Exclude DAB (membrane staining) | FALSE | FALSE | FALSE |
| **Cell parameters** | | | |
| Cell expansion | 1.0 µm | 1.0 µm | 1.0 µm |
| Include cell nulcues | TRUE | TRUE | TRUE |
| **General parameters** | | | |

| | | | |
|---|---|---|---|
| Smooth boundaries | TRUE | TRUE | TRUE |
| Make measurements | TRUE | TRUE | TRUE |
| **Intensity threshold parameters** | | | |
| Score compartment | Nucleus: DAB OD mean | Nucleus: DAB OD mean | Nucleus: DAB OD mean |
| Threshold 1+ | 0.26 | 0.30 | 0.20 |
| Threshold 2+ | (0.4) | (0.4) | (0.4) |
| Threshold 3+ | (0.6) | (0.6) | (0.6) |
| Single threshold | TRUE | TRUE | TRUE |

**Script for the analysis of γH2AX immunohistochemistry staining in LNCaP AR isogenic tumors:**

```
setImageType('BRIGHTFIELD_H_DAB');
setColorDeconvolutionStains('{"Name" : "H-DAB estimated", "Stain 1" : "Hematoxylin", "Values 1" : "0.62644 0.62296 0.46851", "Stain 2" : "DAB", "Values 2" : "0.39554 0.53531 0.74632", "Background" : " 238 239 233"}');
runPlugin('qupath.imagej.detect.tissue.SimpleTissueDetection2', '{"threshold":212,"requestedPixelSizeMicrons":2.0,"minAreaMicrons":10000.0,"maxHoleAreaMicrons":1000.0,"darkBackground":false,"smoothImage":true,"medianCleanup":true,"dilateBoundaries":false,"smoothCoordinates":true,"excludeOnBoundary":false,"singleAnnotation":true}')
selectAnnotations();
runPlugin('qupath.imagej.detect.cells.PositiveCellDetection', '{"detectionImageBrightfield":"Optical density sum","requestedPixelSizeMicrons":0.5,"backgroundRadiusMicrons":85.0,"backgroundByReconstruction":true,"medianRadiusMicrons":0.0,"sigmaMicrons":2.0,"minAreaMicrons":20.0,"maxAreaMicrons":400.0,"threshold":0.26,"maxBackground":10.0,"watershedPostProcess":true,"excludeDAB":false,"cellExpansionMicrons":1.0,"includeNuclei":true,"smoothBoundaries":true,"makeMeasurements":true,"thresholdCompartment":"Nucleus: DAB OD mean","thresholdPositive1":0.26,"thresholdPositive2":0.4,"thresholdPositive3":0.6000000000000001,"singleThreshold":true}')
```

**Script for the analysis of γH2AX immunohistochemistry staining in DLD-1 isogenic tumors:**

```
setImageType('BRIGHTFIELD_H_DAB');
setColorDeconvolutionStains('{"Name" : "H-DAB estimated", "Stain 1" : "Hematoxylin", "Values 1" : "0.63305 0.61773 0.46654", "Stain 2" : "DAB", "Values 2" : "0.36417 0.57223 0.7348", "Background" : " 242 244 239"}');
runPlugin('qupath.imagej.detect.tissue.SimpleTissueDetection2', '{"threshold":212,"requestedPixelSizeMicrons":2.0,"minAreaMicrons":10000.0,"maxHoleAreaMicrons":1000.0,"darkBackground":false,"smoothImage":true,"medianCleanup":true,"dilateBoundaries":false,"smoothCoordinates":true,"excludeOnBoundary":false,"singleAnnotation":true}')
```

```
selectAnnotations();
runPlugin('qupath.imagej.detect.cells.PositiveCellDetection', '{"detectionImageBrightfield":"Optical density sum","requestedPixelSizeMicrons":0.5,"backgroundRadiusMicrons":8.0,"backgroundByReconstruction":false,"medianRadiusMicrons":0.0,"sigmaMicrons":1.4,"minAreaMicrons":10.0,"maxAreaMicrons":400.0,"threshold":0.16,"maxBackground":2.0,"watershedPostProcess":true,"excludeDAB":false,"cellExpansionMicrons":1.0,"includeNuclei":true,"smoothBoundaries":true,"makeMeasurements":true,"thresholdCompartment":"Nucleus: DAB OD mean","thresholdPositive1":0.3,"thresholdPositive2":0.4,"thresholdPositive3":0.6000000000000001,"singleThreshold":true}')
runObjectClassifier("connective tissue DLD-1");
```

**Script for the analysis of γH2AX immunohistochemistry staining in HCT116 isogenic tumors:**

```
setImageType('BRIGHTFIELD_H_DAB');
setColorDeconvolutionStains('{"Name" : "H-DAB estimated", "Stain 1" : "Hematoxylin", "Values 1" : "0.62714 0.62104 0.47012", "Stain 2" : "DAB", "Values 2" : "0.45303 0.56698 0.68796", "Background" : " 244 243 236"}');
runPlugin('qupath.imagej.detect.tissue.SimpleTissueDetection2', '{"threshold":212,"requestedPixelSizeMicrons":2.0,"minAreaMicrons":10000.0,"maxHoleAreaMicrons":1000.0,"darkBackground":false,"smoothImage":true,"medianCleanup":true,"dilateBoundaries":false,"smoothCoordinates":true,"excludeOnBoundary":false,"singleAnnotation":true}')
selectAnnotations();
runPlugin('qupath.imagej.detect.cells.PositiveCellDetection', '{"detectionImageBrightfield":"Optical density sum","requestedPixelSizeMicrons":0.5,"backgroundRadiusMicrons":50.0,"backgroundByReconstruction":false,"medianRadiusMicrons":0.0,"sigmaMicrons":1.3,"minAreaMicrons":13.0,"maxAreaMicrons":400.0,"threshold":0.16,"maxBackground":2.0,"watershedPostProcess":true,"excludeDAB":false,"cellExpansionMicrons":1.0,"includeNuclei":true,"smoothBoundaries":true,"makeMeasurements":true,"thresholdCompartment":"Nucleus: DAB OD mean","thresholdPositive1":0.2,"thresholdPositive2":0.4,"thresholdPositive3":0.6000000000000001,"singleThreshold":true}')
```
